# Antagonism between killer yeast strains as an experimental model for biological nucleation dynamics

**DOI:** 10.1101/2020.09.08.288423

**Authors:** Andrea Giometto, David R Nelson, Andrew W Murray

## Abstract

Antagonistic interactions are widespread in the microbial world and affect microbial evolutionary dynamics. Natural microbial communities often display spatial structure, which affects biological interactions, but much of what we know about microbial warfare comes from laboratory studies of well-mixed communities. To overcome this limitation, we manipulated two killer strains of the budding yeast *Saccharomyces cerevisiae,* expressing different toxins, to independently control the rate at which they released their toxins. We developed mathematical models that predict the experimental dynamics of competition between toxin-producing strains in both well-mixed and spatially structured populations. In both situations, we experimentally verified theory’s prediction that a stronger antagonist can invade a weaker one only if the initial invading population exceeds a critical frequency or size. Finally, we found that toxin-resistant cells and weaker killers arose in spatially structured competitions between toxin-producing strains, suggesting that adaptive evolution can affect the outcome of microbial antagonism in spatial settings.

## Introduction

Microbes affect nearly every aspect of life on Earth, from carbon fixation (Falkowski, Barber, & Smetacek, 1998) to human health (Srivastava & Bhargava, 2016). They often live in dense aggregates, such as biofilms, which offer them protection from environmental forces, drugs and predation (Nadell, Drescher, & Foster, 2016). To prosper in these dense communities and to resist external attacks or takeover from cheater phenotypes, microbes display a wide range of social interactions (West, Diggle, Buckling, Gardner, & Griffins, 2007), both cooperative, such as cross-feeding and quorum sensing, and antagonistic, such as toxin and antibiotic production. The high densities and close proximity of the members of cellular aggregates affect these social interactions, which in turn affect the spatial structure and the spatiotemporal dynamics of microbial communities (Kayser, Schreck, Yu, Gralka, & Hallatschek, 2018).

Laboratory experiments with genetically engineered microbes have helped us understand how social interactions can alter the evolutionary dynamics of microbial populations (Amor, Montanez, Duran-Nebreda, & Sole, 2017; McNally et al., 2017; Muller, Neugeboren, Nelson, & Murray, 2014; Ozgen, Kong, Blanchard, Liu, & Lu, 2018; Weber, Poxleitner, Hebisch, Frey, & Opitz, 2014). For example, cooperation, in which two strains feed each other amino acids, prevents the separation between different genotypes (Muller et al., 2014) that occurs when two non-interacting populations spread across a surface in a range expansion (Hallatschek, Hersen, Ramanathan, & Nelson, 2007). Antagonistic interactions are found in archaea (Atanasova, Pietila, & Oksanen, 2013; Cheung, Danna, O’Connor, Price, & Shand, 1997), prokaryotes (Riley & Wertz, 2002, Veening & Blokesch, 2017) and eukaryotes (Boynton, 2019). They occur in many ecological niches such as the rhizosphere (Kent & Triplett, 2002), aquatic systems (Feichtmayer, Deng, & Griebler, 2017, Drebes Dörr, & Blokesh, 2020) and human infections (Schoustra, Dench, Dali, Aaron, & Kassen, 2012, Libberton, Horsburgh, & Brockhurst, 2015, Heilbronner, Krismer, Brötz-Oesterhelt, & Peschel, 2021), and are frequently exploited for biocontrol applications (Kim et al., 2006; Weller, 2007). These interactions are typically mediated by toxins that are produced and released by cells, e.g. bacteriocins (Schoustra et al., 2012) or antibiotics (Granato, Meiller-Legrand, & Foster, 2019), or injected directly into neighboring cells, as in the case of type VI secretion systems (Borgeaud, Metzger, Scrignari, & Blokesch, 2015, Granato et al., 2019). On an agar plate, the ability of two *Vibrio cholerae* strains to kill each other, using the Type VI secretion system, coarsens the single-strain domains of populations that were initially well-mixed (McNally et al., 2017; Yanni, Marquez-Zacarias, Yunker, & Ratcliff, 2019).

Theoretical models predict different outcomes for cooperation and antagonism: cooperators require each other to prosper (Muller et al., 2014) and antagonistic interactions lead to the competitive exclusion of one of the antagonists (Lavrentovich & Nelson, 2019; Nowak, Sasaki, Taylor, & Fudenberg, 2004; Tanaka, Stone, & Nelson, 2017). We refer to the strain that survives in a 1:1, well-mixed culture as the stronger antagonist and the one that goes extinct as the weaker antagonist. Models based on generalizations of the Lotka-Volterra equations (Lavrentovich & Nelson, 2019, Tanaka, Stone, & Nelson, 2017) predict that being a stronger antagonist is a necessary, but not a sufficient condition for an invading strain to replace a resident, antagonist population: successful replacement requires that the initial inoculum of the invading antagonist be larger than a critical frequency (i.e., relative abundance) in well-mixed populations or a critical size in spatially structured populations. For simplicity, we refer to critical frequency in well mixed populations and critical size in spatially structured ones as ‘critical inoculum size’. The prediction of a critical size has implications for the population dynamics of antagonistic interactions: the requirement for a critical inoculum size implies that mutations conferring an increased strength of antagonism may not necessarily establish in a population because of a deterministic push to extinction if the population size of the mutant is below a given threshold. Conversely, the critical size predicts that populations of weaker antagonists should be resistant to invasion from a stronger antagonist, at least below a certain rate of immigration. Finally, a critical inoculum size has implications for the possibility of exploiting antagonistic, microbial interactions to manipulate microbiomes: exploiting antagonistic interactions may allow us to design microbial consortia that are resistant to external invasion, but if we want to manipulate natural communities, engineered strains will need to be introduced into the microbial community at sufficiently large densities (de Lorenzo, Marlière, & Solé, 2016). The prediction that being a stronger antagonist is necessary, but not sufficient, to invade a resident population requires experimental verification, which motivated our work.

We developed an experimental system in which two strains of the budding yeast, *Saccharomyces cerevisiae*, expressed two different toxins from two different, inducible promoters (Figure 1A). We investigated the dynamics of competitive exclusion in three environments: spatially well-mixed populations in liquid cultures or on surfaces, and spatially structured populations on surfaces. We derived mathematical models of population dynamics regulated by toxin production and toxin-induced cell death and parametrized them using competition assays between toxin-producing (“killer”) cells and sensitive, nonkiller ones (Figure 1B). Experiments verified theoretical predictions on the conditions that lead to a successful invasion of an antagonistic strain in all three environments (Figure 1C). The mathematical models correctly predict the dynamics of competition between toxin-producing strains in all scenarios considered here, they highlight the processes that lead to a region devoid of cells between two antagonistic strains that encounter each other on a solid surface, and can guide attempts to manipulate naturally occurring microbial communities.

**Figure 1.**
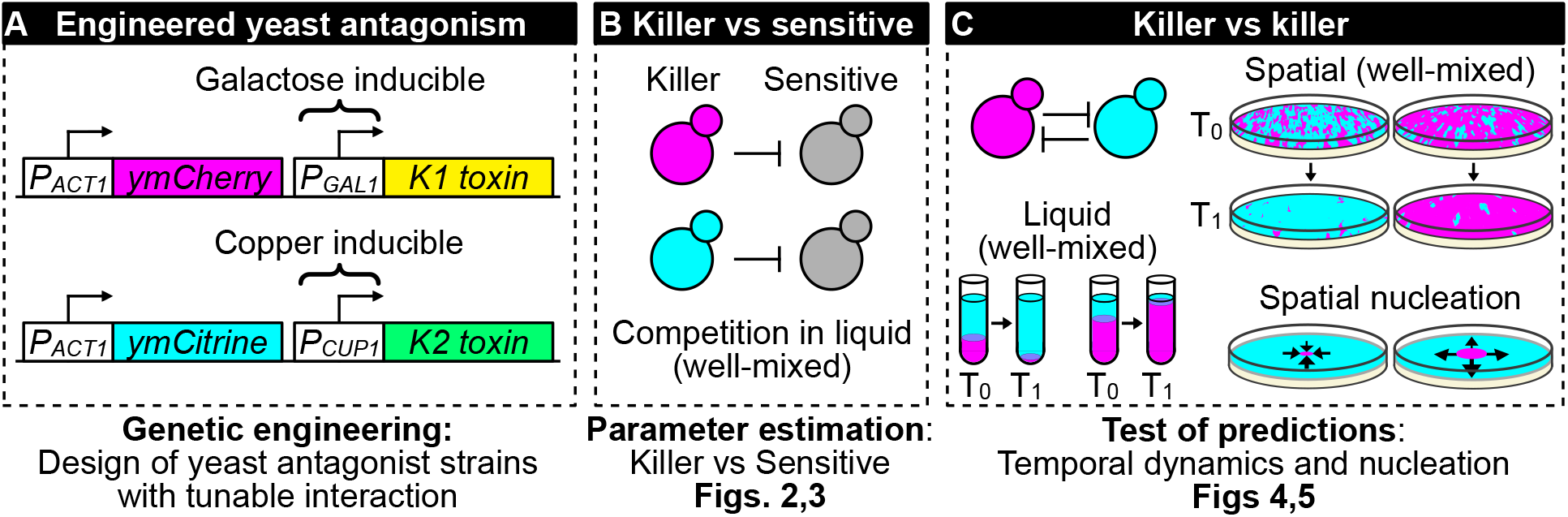
**(A)** We genetically engineered yeast strains to express two different fluorescent proteins (ymCherry and ymCitrine) constitutively and two different toxin/immunity genes (K1 and K2) in response to the inducers galactose (via the *P_GAL1_* promoter) and copper (via the *P_CUP1_* promoter). **(B)** We used competition assays between toxin-producing cells (“killer cells” in cyan and magenta) and sensitive, nonkiller cells (gray cells) to parametrize mathematical models of toxin production, cell growth and toxin-induced cell death. **(C)** We used the models and the experimental system to investigate population dynamics in the presence of antagonistic interactions in both well-mixed (in liquid and on surfaces) and spatially structured populations on surfaces.

We begin by discussing the experimental system (Figure 1A) and the parametrization of mathematical models of antagonism using well-mixed experiments (Figure 1B). Then, we verify the predictions of these models for the competition of two mutually antagonistic strains in three settings: well-mixed cultures in liquid, well-mixed communities on surfaces and spatially structured communities on surfaces (Figure 1C). We then discuss mathematical models that give us intuition for the formation of depletion zones at the interface between two antagonist strains in spatially structured communities and finally we discuss mutants that appeared during the experiments and affected the dynamics of antagonism.

## Results

### Competition between killer and sensitive strains measures toxin production

We began by using competition between killer and non-killer strains in liquid cultures to estimate the parameters needed to model the antagonism between two different killer strains, and to test our ability to vary the strength of the interaction by changing the concentration of the two inducers. The two killer strains expressed two different killer proteins (Tipper & Bostian, 1984) that do not confer immunity to each other: strain K1 expresses the killer toxin K1 (Bevan & Makower, 1963) and strain K2 expresses the killer toxin K2 (Naumova & Naumova, 1973). Killer cells are immune to the toxin they produce because the unprocessed toxins confer immunity (Dignard, Whiteway, Germain, Tessier, & Thomas, 1991; Hanes, Burn, Sturley, Tipper, & Bostian, 1986). Both the K1 and K2 killer toxins bind to β-1,6-glucans on the cell wall, and subsequently translocate to the cytoplasmic membrane where they bind to a secondary receptor (Kre1p for K1, an unknown receptor for K2). Both K1 and K2 disrupt the cytoplasmic membrane increasing its permeability to ions (Magliani, Conti, Gerloni, Bertolotti, & Polonelli, 1997). The nonkiller strains S1 and S2 carried genetic constructs like those in the K1 and K2 strains, but without the killer toxin genes. As a result, S1 expresses the same fluorescent proteins as K1 and S2 expresses the same fluorescent protein as K2, allowing us to distinguish killer cells from nonkiller ones (competing K1 against S2 and K2 against S1), and thus to measure the cell densities of the two strains with a flow cytometer. Figure 2 shows the result of mixing killer and sensitive strains in a 1:1 ratio and following the fraction of the killer strain against time. In the absence of the inducer, galactose, the frequency of K1 remained constant. Increasing the concentration of galactose increased the rate at which the frequency of K1 grew with time. For the K2 toxin, we used two killer strains which had the same genetic construct integrated into their genome, but displayed different fluorescent intensities and different killing strengths (i.e., the rates at which they increased their frequency with time), revealing that the genetic construct was integrated at different copy numbers in the two strains, K2 and K2_b_. For both strains, their relative frequency increased with time even in the absence of copper, which is consistent with leaky expression from the *P_CUP1_* promoter (Butt et al., 1984; Gorman, Clark, Lee, Debouck, & Rosenberg, 1986) in the absence of added copper. Increasing the concentration of copper increased the rate at which the two strains’ frequencies grew with time; the less fluorescent strain K2_b_ was a weaker killer than the more fluorescent strain K2 at all copper concentrations.

**Figure 2.**
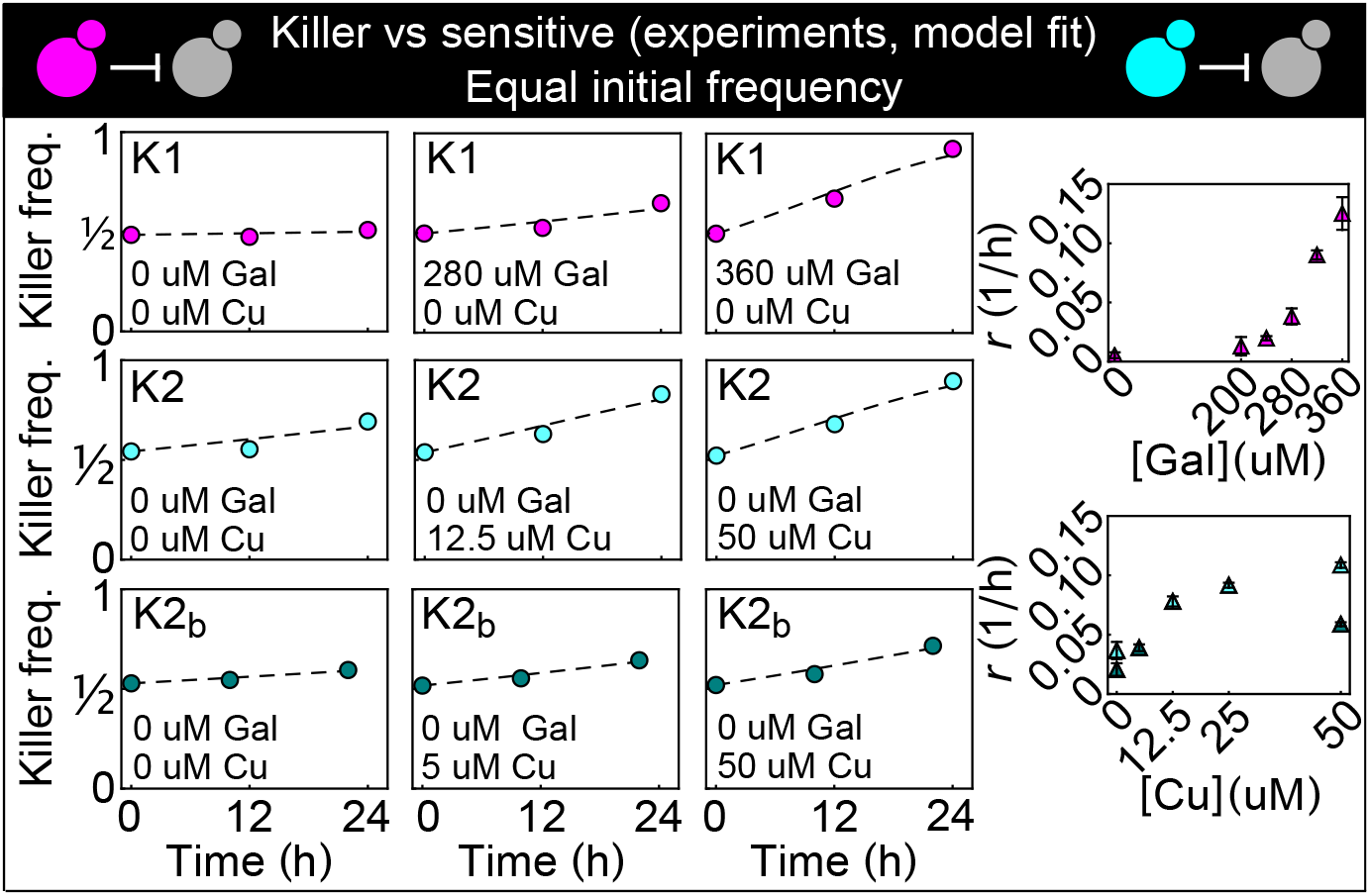
Temporal change of the killer strain frequencies in competition assays against sensitive strains, at different inducer concentrations. Different colored points depict data corresponding to the killer strains K1 (magenta), K2 (cyan) and K2_b_ (dark green). For each killer strain, increasing its inducer’s concentration increased its killing strength, that is, the rate at which its frequency grew with time. In the absence of inducers, the K1 strain frequency remained constant, whereas strains K2 and K2_b_ still displayed killing activity, which we attribute to the leakiness of the *P_CUP1_* promoter. Each data point is the mean of two, three or five technical replicates. The x axis reports time since the first measurement. Dashed lines show the best fits of the frequency model (Equations 1 and 10). The panels on the right show the best-fit interaction coefficients (which are proportional to toxin production rates) as a function of the inducer concentrations (mean±SD, Table 3).

We developed a simple mathematical model of cell growth, toxin production and toxin-induced cell death and used the data in Figure 2 to fix the parameters. Under suitable assumptions on the relative time scales of cell division and toxin production (Methods), the temporal change of the K1’s frequency, *f,* in competition against a K2 killer strain under well-mixed conditions can be described by a single equation:

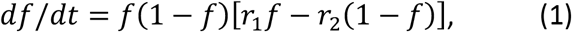

where *df/dt* denotes the temporal derivative of *f*, and *r*_1_ and *r*_2_ are interaction coefficients proportional to the toxin production rates of strains K1 and K2 (see Methods). For two strains K1 and K2 at equal initial frequencies (i.e., *f*_0_ = 1/2), a Taylor-expansion of the solution to Equation (1) around *f*_0_ shows that, initially, *f* varies linearly with time with a rate proportional to *r*_1_ − *r*_2_, i.e. 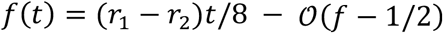, before non-linear terms become important. For a killer strain competing against a sensitive, nonkiller one, Equation 1 reduces to *df /dt* = *rf*^2^(1 − *f*), where *f* is the killer strain frequency. The best fits of this last equation are shown as dashed lines in Figure 2, and the best-fit estimates of the interaction coefficient *r* for the three strains K1, K2 and K2_b_ at different inducer concentrations are shown in the lower panels (numerical values are given in Table 3). A formally equivalent model that included spatial diffusion and noise due to number fluctuations was studied theoretically in (Lavrentovich & Nelson, 2019), where it was derived starting from a stepping-stone model with local, antagonistic interactions, i.e. without explicitly modeling the secretion of diffusible toxins. When toxin dynamics are much faster than cell density dynamics, our model and the well-mixed version of the earlier model (Lavrentovich & Nelson, 2019) coincide (Methods).

**Table 1.**
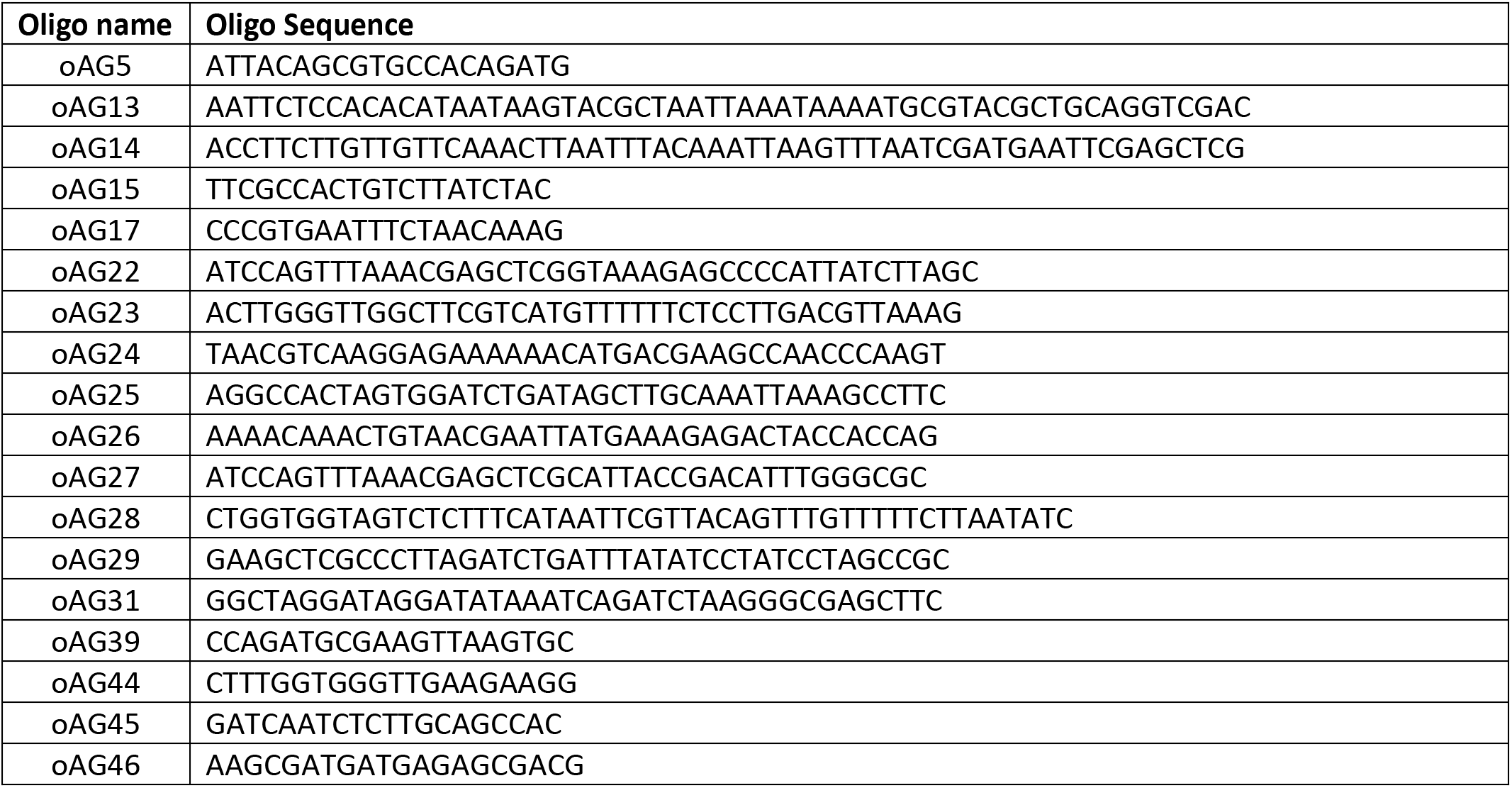
Oligos used in this study.

**Table 2.**
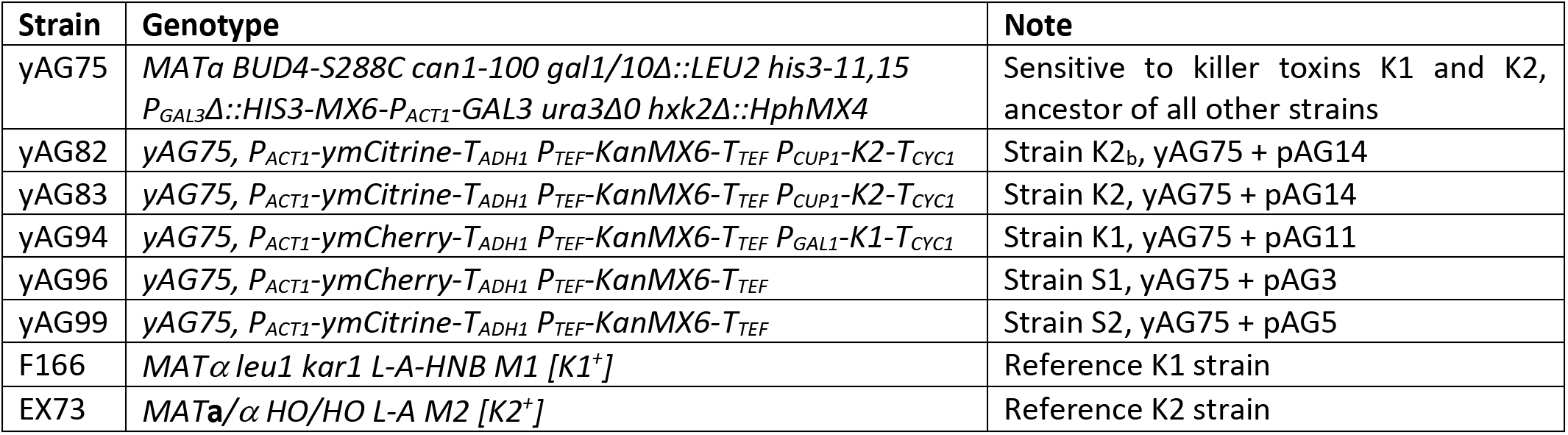
Strains used in this study. All strains are in the *S. cerevisiae* W303 background. Strain yAG75 has not been used in the experiments but is reported here because it’s the ancestor of all the other strains; only those elements that differ from yAG75 are listed for the other yAG strains.

**Table 3.** Best-fit estimates for the parameters of the frequency model Equation (10) fitted to the data from competition assays between toxin-producing strains and sensitive, nonkiller ones. Concentrations of the inducers galactose (Gal) and copper (Cu) are indicated as superscripts, whereas the parameters *r*_1_, *r*_2_ and *r*_2*b*_ (i.e., the parameter *r* in Equation 10 for the strains K1, K2 and K2_b_) are given in units of 1/h (mean±SD).

|  |  |  |  |  |  |  |
| --- | --- | --- | --- | --- | --- | --- |
| $r_1^{0 \text{ uM Gal}}$ | $r_1^{200 \text{ uM Gal}}$ | $r_1^{240 \text{ uM Gal}}$ | $r_1^{280 \text{ uM Gal}}$ | $r_1^{320 \text{ uM Gal}}$ | $r_1^{360 \text{ uM Gal}}$ | |
| $0.005 \pm 0.002$ | $0.013 \pm 0.008$ | $0.020 \pm 0.002$ | $0.038 \pm 0.007$ | $0.090 \pm 0.004$ | $0.125 \pm 0.014$ | |
| $r_2^{0 \text{ uM Cu}}$ | $r_2^{12.5 \text{ uM Cu}}$ | $r_2^{25 \text{ uM Cu}}$ | $r_2^{50 \text{ uM Cu}}$ | $r_{2b}^{0 \text{ uM Cu}}$ | $r_{2b}^{5 \text{ uM Cu}}$ | $r_{2b}^{50 \text{ uM Cu}}$ |
| $0.037 \pm 0.007$ | $0.078 \pm 0.004$ | $0.092 \pm 0.002$ | $0.109 \pm 0.002$ | $0.021 \pm 0.005$ | $0.039 \pm 0.003$ | $0.059 \pm 0.002$ |

According to Equation 1, the dynamical system describing the antagonistic interaction of two killer strains has two stable equilibria, one at *f* = 0 and one at *f* = 1, and one unstable equilibrium at *f_eq_* = *r*_2_/(*r*_1_ + *r*_2_). If the initial frequency is above *f_eq_* the system tends to *f* = 1, otherwise it tends to *f* = 0. In other words, the strain K1 can only increase its frequency in the population if its initial frequency is larger than the critical inoculum frequency, *f_eq_*. The equilibrium frequency *f_eq_* thus represents a critical inoculum size below which the invasion of a stronger antagonist is predicted to fail in well-mixed settings. Note that this particular “size” relates to an inoculum *concentration* rather than the actual physical size discussed later in this paper for spatially structured communities on surfaces. Nevertheless, when number fluctuations are included in the dynamics, there is an interesting analogy with escape over a barrier problems in statistical mechanics (Chotibut & Nelson, 2015). Increasing the toxin production rate of strain 1 increases *r*_1_, and is thus predicted to decrease the size of the critical inoculum. An intuitive derivation for the critical frequency *f_eq_* can be obtained by assuming that the two toxins have equal per-cell binding rates and kill cells of the other strain at the same rate: With this assumption, *r*_1_ and *r*_2_ are proportional to the per-cell toxin production rate of the strains K1 and K2, and the equilibrium frequency *f_eq_* is the frequency at which the populations of the two strains produce the same amount of toxins per unit time. In the general case in which the two toxins have different per-cell binding and killing rates, the equilibrium frequency *f_eq_* is such that the toxin-induced, per-capita death rates for the two strains are equal. A critical inoculum size is thus present in this system because a stronger antagonist must overcome the toxin production from its competitor, before being able to expand in the population. Figure 3C plots the rate at which the fraction of strain 1 changes at different interaction coefficients, which are determined by the concentrations of galactose and copper, the inducers of toxin production. Equation 1 can also be rewritten as *df /dt* = −*dV/df*, where *V* is the quartic potential depicted in Figure 3C. When both *r*_1_ and *r*_2_ are positive, the potential *V* has a double-well structure with two minima, corresponding to the two stable states in which one strain competitively excludes the other. Separating the two minima is an energy barrier having its peak at the critical frequency that the first strain must overcome to exclude the second. As shown theoretically in (Lavrentovich & Nelson, 2019), the spatial, stochastic generalization of Equation 1 can be interpreted as an escape over the barrier problem, and lends itself to the use of theoretical techniques from nucleation theory as first appreciated by Rouhani and Barton (Rouhani & Barton, 1987) in the context of spatial population genetics.

**Figure 3.**
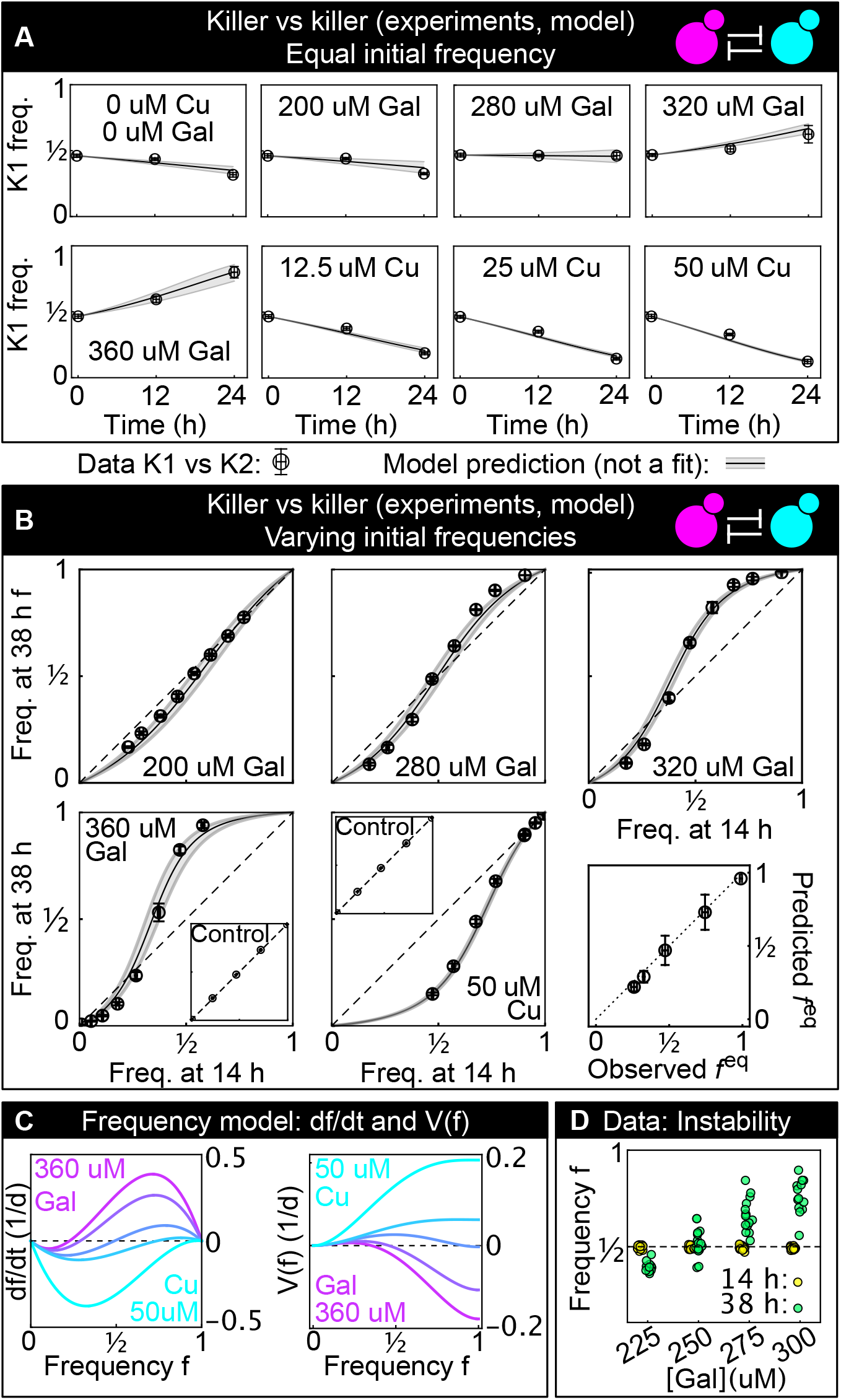
**(A)** Temporal change of strain K1’s frequency in competition assays against strain K2, at different inducer concentrations (only one of the two inducers was added in each replicate). Each data point is the mean of 13 or more technical replicates, error bars are one standard deviation. Solid lines in (A) and (B) are predictions according to Equation 1 with parameters estimated from competitions between the killer strains K1 and K2 and the sensitive, nonkiller strains S1 and S2. Gray bands show the 68% confidence interval for the model. **(B)** Changes in the frequency of strain K1 following 24 h of competition against strain K2, at different concentrations of the inducers (subpanels) and at different initial frequencies. The x axis gives the K1 frequency 14 h after inoculation, the y axis gives the K1 frequency 38 h after inoculation. The dashed lines show the 1:1 line that points would lie on if the two strains had equal fitness. The critical inoculum corresponds to the intersection point between the dashed lines and the solid lines (model), or an interpolation of the data points (experiment). Insets show the control experiments of competing the two sensitive strains S1 and S2 with each other at the same inducer concentrations as the parent subpanels. The bottom-right subpanel shows the correlation between the value of *f_eq_* predicted from our model (the intersection point between the solid and dashed lines) and the experimental value of *f_eq_* based on competitions (the intersection of the interpolation between the data points and the dashed lines) for all inducer concentrations. The dotted line is the 1:1 line. **(C)** Temporal derivative of *f* according to Equation 1, at the inducer concentration values of panel B (left, only the lines corresponding to the highest galactose and highest copper concentrations are labeled) and the corresponding quartic potential (right). **(D)** At the galactose concentration of 250 uM, the unstable equilibrium *f_eq_* is close to ½. Different technical replicates that start around *f* = ½ tend towards different stable equilibria of Equation (1) (i.e. *f* = 0 and *f* = 1) in the long-term limit, highlighting the instability of the equilibrium point. Yellow points show frequencies of the K1 strain 14 h after inoculation, green points show frequencies of the K1 strain 38 h after inoculation.

### A simple model predicts the competition between two antagonistic killer strains

Having measured the two interaction coefficients (*r*_1_ and *r*_2_) in competitions involving antagonism acting on sensitive strains, we asked if our model could predict the frequency dynamics of the two killer strains K1 and K2 competing against each other. Figure 3A shows the frequencies of the K1 and K2 strains grown in liquid following the same protocol as the experiments of Figure 2, starting from equal frequencies for the two strains and varying the concentrations of the inducers, which control *r*_1_ and *r*_2_. The frequency of K1 increased if *r*_1_ > *r*_2_ and decreased otherwise, in accordance with Equation 1 above. Figure 3B shows the frequencies of the two strains separated by an interval of 24 h, with the frequencies at 14 h after inoculation on the x axis and the frequencies at 38 h after inoculation on the y axis. The insets in panel B show two control experiments consisting of competition assays between the nonkiller strains S1 and S2: the two strains have identical fitness, so their relative frequency remains constant over 24 h of growth. The model predictions (solid line) and the 68% confidence intervals (gray shading) reveal that the model can predict the temporal dynamics and the value of the unstable equilibrium *f_eq_* (last panel in Figure 3B), for all inducer concentrations and for all initial frequencies. The ability of parameters estimated from killer vs sensitive assays (Figure 2) to predict the dynamics of the competition between two killer strains shows that the interaction terms included in Equation 1 are sufficient to capture the antagonistic dynamics: we can simply sum the contribution of strain K2 to the death rate of strain K1 (the term *r*_2_ (1 − *f*) in Equation 1, see Methods) and the corresponding contribution of strain K1 to the death rate of strain K2 (the term *r*_1_*f*) without adding additional terms to the equation. The experiments also show that for initial frequencies close to the unstable equilibrium, different technical replicates can tend towards different stable equilibria in the long-term limit (Figure 3D), highlighting the instability of the equilibrium *f_eq_* and the fact that the energy barrier can be overcome when the initial frequency is close to *f_eq_*. Figure 3 - supplement 2 shows that there is no correlation between the frequency at the first and second measurement time point for those replicates that were initialized close to the unstable equilibrium (i.e., data points at 250 uM galactose in Figure 3D), suggesting that the stochasticity of the dynamics dominates over the initial condition when determining which replicates tend towards different stable equilibria in the long-term limit. Overall, the experimental results from well-mixed experiments (Figure 3) confirm the theoretical prediction that a critical starting frequency, the equilibrium frequency *f_eq_*, is required for a stronger antagonist to invade a resident, antagonist population.

Because many microbial communities are spatially structured, we studied the spatial dynamics of antagonism in *spatially* structured populations growing on surfaces. As an intermediate step between well-mixed liquid cultures and spatially structured populations on surfaces, we studied the interaction between antagonistic strains on a solid surface; we distributed an initially well-mixed population of the two killer strains K1 and K2 on the surface of agar plates, with different concentrations of the inducers and with different initial frequencies of the two strains (Figure 4). We let the two strains grow for 24 h and then measured their relative frequencies using a fluorescence stereomicroscope. These experiments were designed to investigate if the same inducer concentrations used in liquid led to similar strengths of antagonism between the two toxin-producing strains growing on the surface of agar plates. We found that increasing the concentration of the two inducers led to increased killing activity for the two strains, and that the copper-induced killer K2 appeared to be a more effective killer on plates than in liquid, as suggested by the fact that the equilibrium points *f_eq_* in liquid (Figure 3B) are smaller than those on plates at comparable concentrations of galactose (Figure 4), and by the observation that the competitive exclusion of K1 by K2 on plates with 12.5 uM copper happened much faster than in liquid with 50 uM copper (compare Figure 4, showing data after 24 h from inoculation on plates, with Figure 3B, showing data after 38 h from inoculation in liquid). Although we do not know why the K2 killer strain was a stronger antagonist on plates than in liquid, one possibility is that the agar (the only ingredient that differs between the liquid and solid media) contained traces of copper (Debergh, 1983) leading to a stronger expression of the K2 toxin genes.

**Figure 4.**
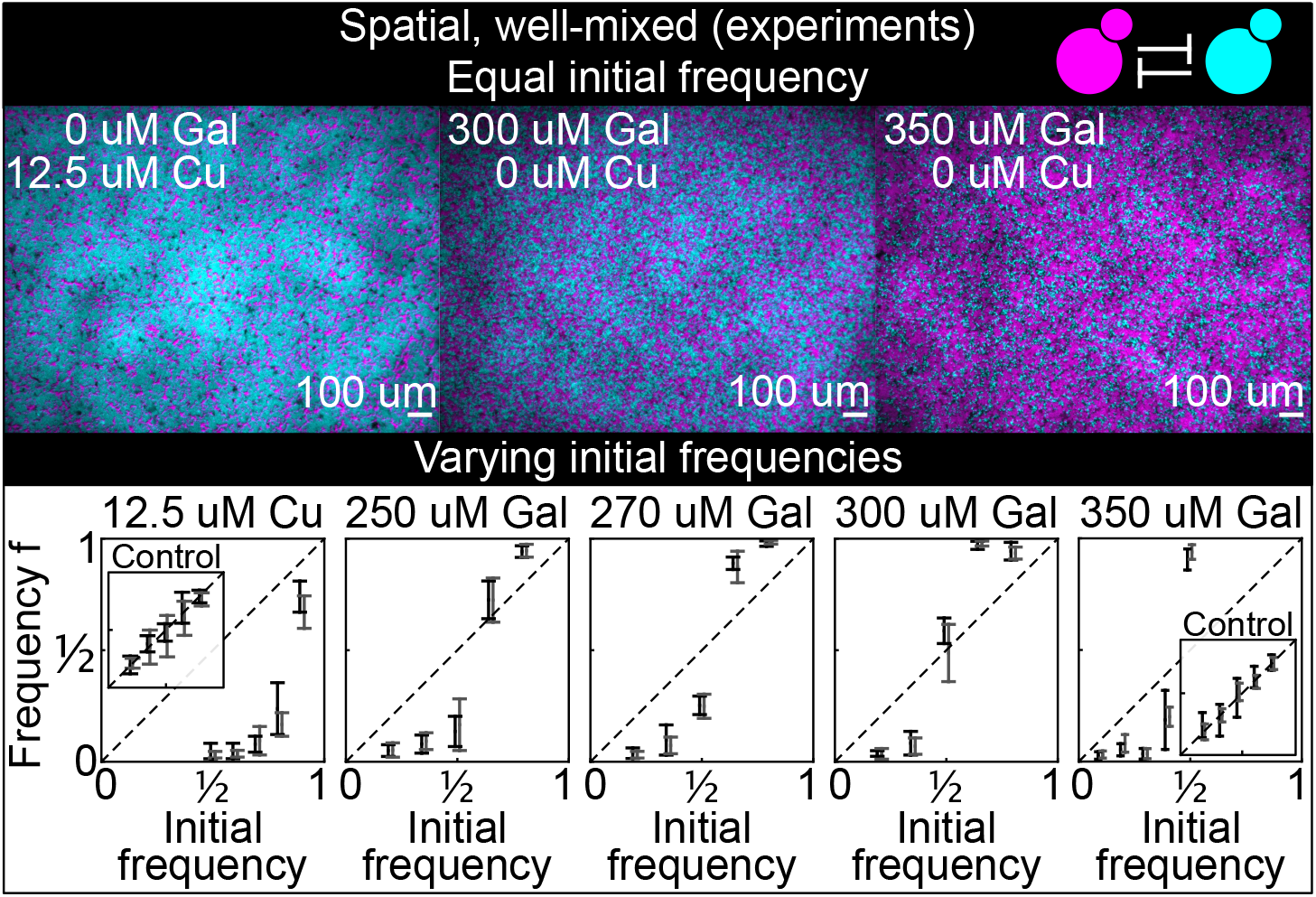
Antagonistic competition of spatially well-mixed, toxin-producing strains K1 and K2 growing on solid agar surfaces, at different inducer concentrations (subpanels) and different initial frequencies. The upper panels show combined fluorescence images of three representative spatially well-mixed populations imaged 5 h after inoculation. The images show populations that started at 50:50 initial frequencies of K1 and K2, with K2 outcompeting K1 on the left and the opposite outcome on the right. The lower panels show the relative frequencies of the two strains at the time of inoculation (x axis) and 24 h after inoculation (y axis), estimated as the relative fraction of space occupied by each strain. Black and gray data show data from two different experiments, and the two whiskers of each data point connect the maximum and the minimum estimated frequency *f* of K1. Due to the difficulty of unequivocally assigning each pixel to one or the other strain in this assay, we report conservative estimates of the maximum and minimum frequencies that we can confidently assign to the K1 strain. The dashed lines show the 1:1 line that points would lie on if the two strains had equal fitness. Insets in the lower sub-panels show control experiments: competition assays between the nonkiller strains S1 and S2, at the same inducer concentrations as the parent subpanels.

### Invasion in spatially structured populations requires a critical inoculum size

Finally, we investigated antagonism dynamics in spatially structured populations. We asked whether an invading antagonist inoculated at one location could invade a surface uniformly occupied by a resident, weaker antagonist (see sketch in Figure 1C). We spread a uniform lawn of the weaker killer strain K2_b_ on the surface of an agar plate and inoculated droplets of different volumes of a culture of strain K1. Experiments were performed with 360 uM galactose, an inducer concentration at which strain K1 is a stronger antagonist than strain K2_b_. We let the two strains grow on the surface of these agar plates for 48 h, following which a fraction of the populations was transferred onto fresh plates by replica plating, providing fresh nutrients while preserving their spatial structure (Figure 5A), and we allowed them to grow for further 48 h. This procedure was repeated for 13 serial transfers. We imaged the populations at the end of each growth period (Figure 5B) using a fluorescence stereomicroscope, and measured the area occupied by the invading strain in each population (Figure 5C-D). As shown in Figure 5B-D, all K1 populations whose initial area was below 12 mm^2^ were outcompeted by the K2_b_ lawn and driven to extinction, whereas all those whose initial area was above 12 mm^2^ managed to persist and eventually expanded displacing the resident K2_b_ population (except for one outlier population highlighted in Figure 5B-D with a gray arrow and gray lines, which we discuss separately below). Most of the populations exhibited two characteristic phases in their spatial dynamics (Figure 5C-E): an initial retreat of both strains, leaving a region devoid of cells at the outer edge of the initial inoculum (black “halos” in Figure 5E, of width 400+200 um, mean+SD), followed by the expansion of the K1 strain. All the inoculations below the critical inoculum size, instead, went extinct very rapidly (Figure 5A,F). A control experiment performed in parallel using the two sensitive, nonkiller strains S1 and S2 shows that all inoculations maintained their area following successive transfers (Figure 5 - supplement 2). Thus, the retreat and expansion dynamics and the dependence of invasion success on the size of the initial inoculum can be attributed to the antagonistic interactions engineered between strains K1 and K2_b_.

**Figure 5.**
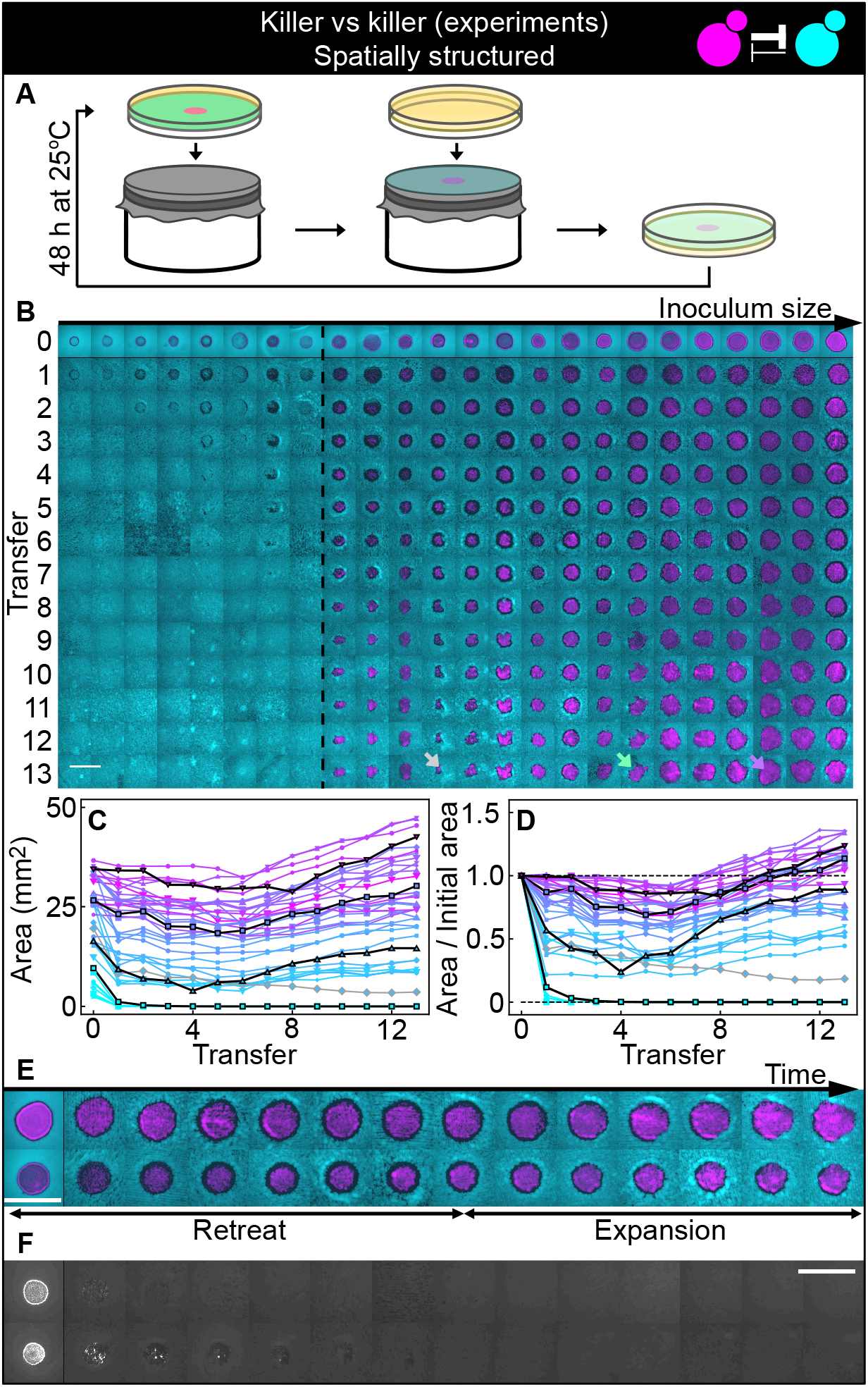
Antagonistic competition of killer strains K1 and K2_b_ in spatially structured populations. Experiments were performed on agar medium with 360 uM galactose, a concentration at which the K1 strain is a stronger killer than the K2_b_ strain. **(A)** We used replica plating to replenish nutrients and dilute the populations, while preserving the spatial structure of the population. At the end of each growth period (48 h at 25°C), the agar plates hosting the experimental populations were gently pressed onto a microfiber cloth laid flat on a cylinder, leaving a diluted copy of the population on the cloth. A fresh agar plate was then pressed onto the same cloth, leading to an effective dilution of the population that preserved its spatial structure. **(B)** Shown from left to right are spatially structured populations of the invader K1 (magenta) and the resident K2_b_ (cyan) strains originated from depositing droplets of different volumes of K1 onto a lawn of K2_b_ cells. The populations are ordered based on the number of K1 cells at the end of the first growth cycle (estimated from the integrated ymCherry fluorescence of strain K1): the population with the smallest number of K1 cells is on the left, that with the largest number on the far right (first row). Different rows in the same column show the same population 48 h after the previous transfer. The existence of a critical inoculum size is clearly visible and is marked by a dashed, black line. Populations on the left of this line failed to expand, whereas populations on the right of it persisted and eventually expanded. (B) shows only a subset of the experimental populations; Figure 5 - figure supplement 1 shows all the populations. An outlier population (gray arrow) and a population with a re-invading K2_b_ subpopulation (green arrow) are discussed in the main text. **(C)** Area covered by each K1 population at the end of each 48-h growth period between transfers, color coded from cyan to magenta according to the integrated ymCherry fluorescence intensity of each replica at the end of the first growth period. Highlighted in black are four characteristic curves highlighting the fact that populations above the critical inoculum initially decrease in size, before expanding later. **(D)** Same data as in (C), divided by the initial area to highlight relative changes. Shown in **(E)** are two populations in which the retreat and expansion phases of the dynamics are clearly visible. The two rows show two different populations, whereas different columns show the same populations at the end of the growth periods following successive transfers. Both K1 (magenta) and K2_b_ (cyan) populations initially retreat, leaving a region without cells (a black “halo” surrounding the magenta islands). In the expansion phase of the dynamics, the magenta regions expand and increase their area. These populations are well above the critical inoculum size. Panel **(F)** shows the temporal dynamics of the two largest populations below the critical inoculum (last two columns of panel B before the dashed, black line). The largest population (first row in F) disappeared almost immediately, whereas the second-to-last one (second row) disappeared after five transfers. Only the fluorescence due to strain K1 is shown. Scale bars are 1 cm long. Figure 5 - supplement 2 shows a control experiment in which we followed the same protocol using the sensitive, nonkiller strains S1 and S2, and found that the areas of S1 inoculations on a lawn of S2 cells remained constant with time, for all initial inoculation sizes.

### Mutations in killer production and sensitivity alter the outcome of competitions

Our theory ignores the possibility that mutations arise which alter the interaction between the antagonistic strains. Some replicates showed dynamics that differed from the typical dynamics described in the previous paragraph suggesting that such mutations occurred at detectable frequency. An outlier K1 population (K1^o^) highlighted with a gray arrow and gray lines in Figure 5B-D (first row of Figure 6A) decreased its area monotonically with time, unlike all other populations that either went extinct or eventually expanded their area. At the end of this experiment, we collected cells from all the experimental populations and made glycerol stocks. Flow cytometry measurements of the fluorescent intensity of K1 cells sampled from the outlier replica K1^o^ showed that their average fluorescent intensity was reduced with respect to the ancestral K1 population, and also compared to K1 cells sampled from a non-outlier population (K1^s^) that successfully expanded in the spatially structured experiments (Figure 6B). Fluorescence imaging of colonies grown from the sampled and ancestor populations confirmed this observation. K1, K1^s^ and K1^o^ all contained both high and low-fluorescence cells, but K1^o^ contained many more low-fluorescence ones than K1 and K1^s^. Because the gene expressing the fluorescent protein ymCherry is close to the K1 toxin gene on our genetic construct (Figure 1A), a hypothesis for the monotonic decrease in the area of the outlier population is that it was on average a weaker antagonist, expressing the K1 killer toxin gene to a reduced degree compared to the ancestor population and other populations that successfully expanded. Such a reduced expression may have occurred due to one of two sorts of mutation: a mutation that greatly elevated the rate of recombination between multiple, tandemly integrated copies of the construct, or a mutation that led to its reversible and epigenetic silencing.

**Figure 6.**
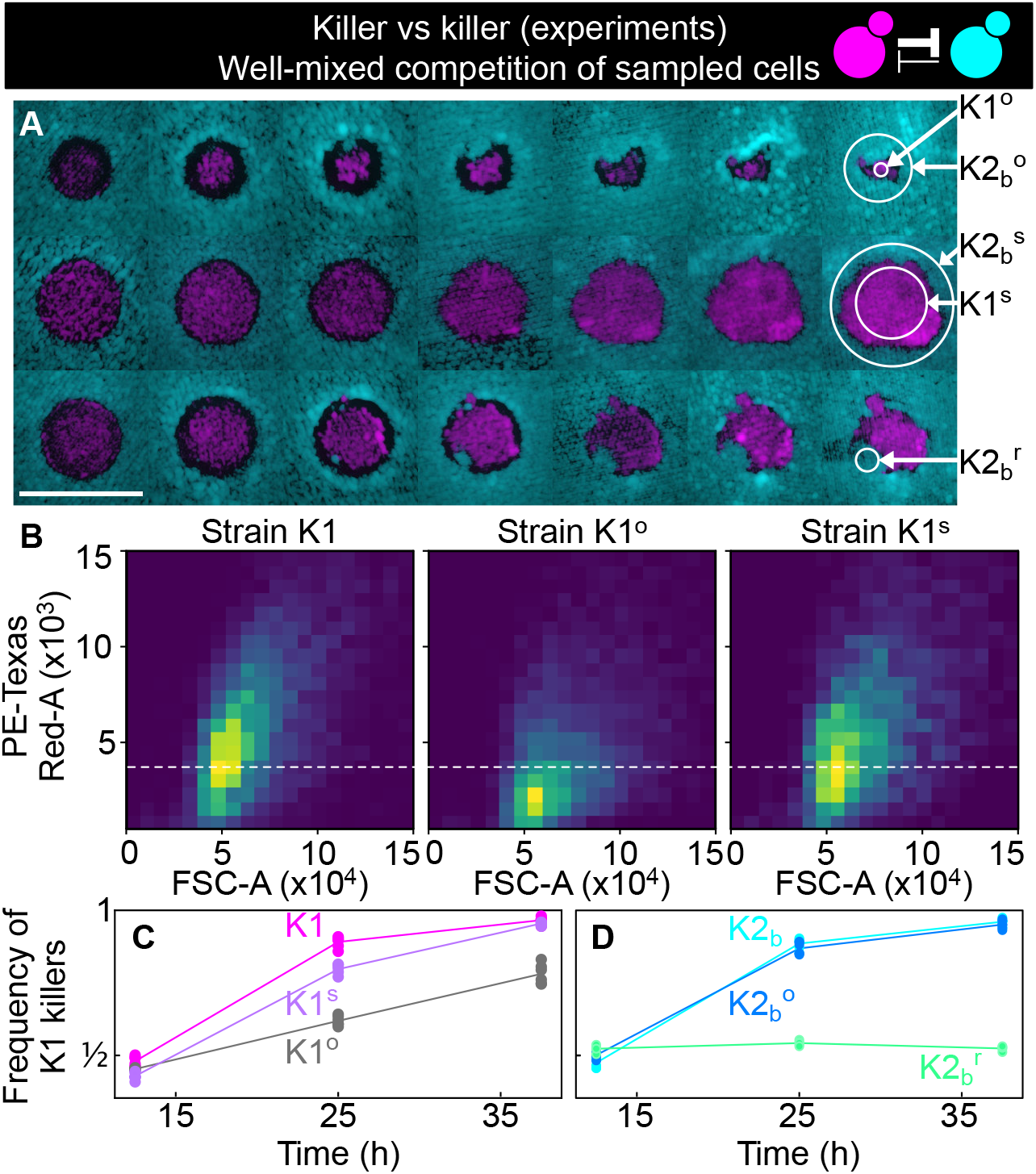
Cells sampled from certain replicates at the end of the experiment shown in Figure 5 showed altered killing strength and toxin-resistance. **(A)** Regions from which the samples were taken at the end of the experiments of Figure 5. The three rows show the replicates highlighted with gray, purple and green arrows in Figure 5B, respectively. Different columns show the same populations at every other transfer. The scale bar is 1 cm long. **(B)** Density histograms of fluorescence intensity (y axis, arbitrary units) vs forward scatter (FSC, x axis, arbitrary units), which correlates with cell size, measured via flow cytometry during competitions in well-mixed liquid cultures between strains K1, K1^o^ and K1^s^, versus K2_b_. K1 is the strain used in all other experiments, K1^o^ and K1^s^ are populations sampled at end of the experiments of Figure 5. The fluorescence intensity of population K1^o^ is lower than that of K1 and K1^s^. The dashed line shows the K1 histogram mode as a visual aid to compare fluorescent intensities. **(C)** In competition assays in liquid at 360 uM galactose, the frequency of K1 and K1^s^ competing against strain K2_b_ increases faster than the frequency of K1^o^ competing against K2_b_, showing that population K1^o^ is a weaker killer than K1 and of other populations that successfully expanded in the experiments of Figure 5 (e.g., K1^s^). **(D)** In competition assays in liquid at 360 um galactose, strain K1 competing against strain K2_b_ (cyan) and strain K1 against the sub-population K2_b_^o^ (light blue) sampled at the end of the experiment of Figure 5 follow similar dynamics suggesting that the collapse of K1^o^ was not due to increased toxin production by K2_b_^o^, or it developing resistance to the K1 toxin. The competition assay with strain K1 against strain K2_b_^r^ (green), which re-invaded a K1 population in the experiments of Figure 5, instead, showed no increase in frequency for strain K1, suggesting that K2_b_^r^ developed resistance to the K1 toxin. Different data points in (C-D) show different technical replicates.

We tested the hypothesis that the outlier population K1^o^ was a weaker antagonist by performing a competition assay in liquid medium, competing K1^o^ against the ancestor K2_b_ population and against the nonkiller strain S2, at 50:50 initial frequencies following the same protocol as the experiments of Figure 2 and Figure 3A. In the same experiment, we competed the ancestor K1 and the successful invader, K1^s^, against the ancestor K2_b_ (Figure 6C and Figure 6 - figure supplement 1, panel A) and against the nonkiller strain S2. We found that K1^s^ and the ancestor K1 increased in frequency faster than K1^o^, demonstrating that the outlier population K1^o^ was indeed a weaker antagonist than K1^s^ and K1, which followed similar trajectories. An alternative hypothesis for the outlier behavior of K1^o^ would be that K2_b_ cells in that region developed resistance to the K1 toxin. Even though visual inspection of the spatiotemporal dynamics of K2_b_ in the outlier replica suggest that this might have happened (cyan cells rapidly re-invaded the magenta K1^o^ population from the top, see Figure 6A), competition assays between K1 and K2_b_ cells sampled from the area surrounding K1^o^ (K2_b_^o^) rule out this hypothesis, given that they followed the same dynamics of competition assays between the original K2_b_ stock and K1 (Figure 6D and Figure 6 - figure supplement 1B). Given the small size of the K2_b_ front that invaded the K1 population from the top (Figure 6A), however, it is possible that when sampling K2_b_^o^ cells from the region surrounding K1^o^ we failed to isolate and test potentially toxin-resistant cells from the invading region. Other populations showed interesting phenotypes, like the one highlighted with a green arrow in Figure 5B (third row of Figure 6A). In this population, K2_b_ cells (cyan) re-invaded the K1 population (magenta) after the halo had formed. By competing K2_b_ cells from the sub-population highlighted with the green arrow (K2_b_^g^) in Figure 5B against the ancestor K1 in competition assays in liquid media with 360 uM galactose, we found that they had greatly decreased (possibly null) sensitivity to the toxin produced by K1 (Figure 6D), suggesting that they became K1-resistant during the course of the experiment. Figure 6 - figure supplements 2 and 3 reveal that we could not detect any differences in growth rate between any pairs of strains, ruling out the possibility that the altered outcomes of competition observed in Figure 6 could be due to changes in cell division times during the experiment.

### Models suggest that nutrient depletion produces unoccupied halos between antagonistic strains

The frequency model used for well-mixed populations cannot be directly applied to describe the experiments with spatially structured populations on surfaces, since it does not include the diffusion of the two toxins, which likely underlies features such as the halo region at the boundary between two antagonistic strains (Figure 5E). To mathematically investigate the spatial dynamics of the two antagonistic strains, we explored a suite of models in which we explicitly modeled the density of each strain (rather than their relative frequency as in Equation 1) as a function of space and time, along with the concentration of the two toxins. We used the parameters for toxin production derived from competition between killers and nonkiller strains (data shown in Figure 2) and estimated the diffusion rates of the toxins and yeast cells based on their size (Table 6). By exploring models with different levels of complexity and realism (Methods), we found that we had to explicitly model the dynamics of nutrients (glucose) to reproduce the formation of the halo, which in the models consists of a region of mutual destruction, with significantly reduced cell density at the boundary between the two strains. In the most realistic model we investigated (Methods), cells occupy a two-dimensional surface at the top of the agar plate and their growth dynamics is modeled via a growth rate that depends on nutrient concentration and saturates with an effective half-saturation constant, Km, for the local nutrient concentration, and a death term proportional to the local concentration of the toxins. Cells diffuse locally on the surface of the agar via a growth-dependent diffusion term reflecting the fact that cells push each other around as they interact mechanically with other cells during their growth and division (Giometto, Nelson, & Murray, 2018; Kayser, Schreck, Gralka, Fusco, & Hallatschek, 2019). The toxins and the nutrients diffuse in the agar and are produced and/or depleted by cells at the surface, at rates that depend on the local density of the two strains. We found that this model, parametrized using the competition assays between toxin-producing and nonkiller strains and using suitable estimates for the diffusion coefficients of the nutrients, toxins and yeast cells taken from the literature, could reproduce the striking halos observed at the boundary between the two strains (Figure 7A), suggesting that the halo emerges due to a combination of toxin-induced killing and diffusion of nutrients away from the agar beneath the halo and their consumption by the cells bordering the halo. Upon numerically integrating the model, starting from initial conditions comparable to those used in the experiments with spatially structured populations, we found that it correctly predicts the extinction of the smaller inoculations, with a critical inoculum size between 8 and 14 mm^2^, and the initial retreat and subsequent expansion of the larger ones (Figure 7A-C). The model also predicts an enhanced density of the two killer strains at the two sides of the halo (Figure 7A), caused by the diffusion of glucose away from the agar underneath the halo to sustain cell growth at the exposed edges of the regions occupied by the two strains. The enhanced density in those regions was seen also in the experiments as an increased fluorescence signal (see Figure 5E), even though the magnitude of this phenomenon appears larger in the experiments than in the model. If the halo was caused by the presence of the toxins alone, and not by the combined effect of the toxins and the diffusion of nutrients away from the agar underneath the halo, one would expect that inhibition of the toxin would allow cells to re-invade the halo region. To test this, we experimentally verified that no further growth in the halo region is observed after transferring populations that competed for 48h at 25°C to 32°C for further 48h (Figure 5 - supplement 3), a temperature at which both the K1 and K2 toxins are unstable and fail to inhibit the growth of susceptible strains (Marquina, Santos, & Peinado, 2002, Lukša, Serva, & Serviene, 2016, Figure - supplement 4).

**Figure 7.**
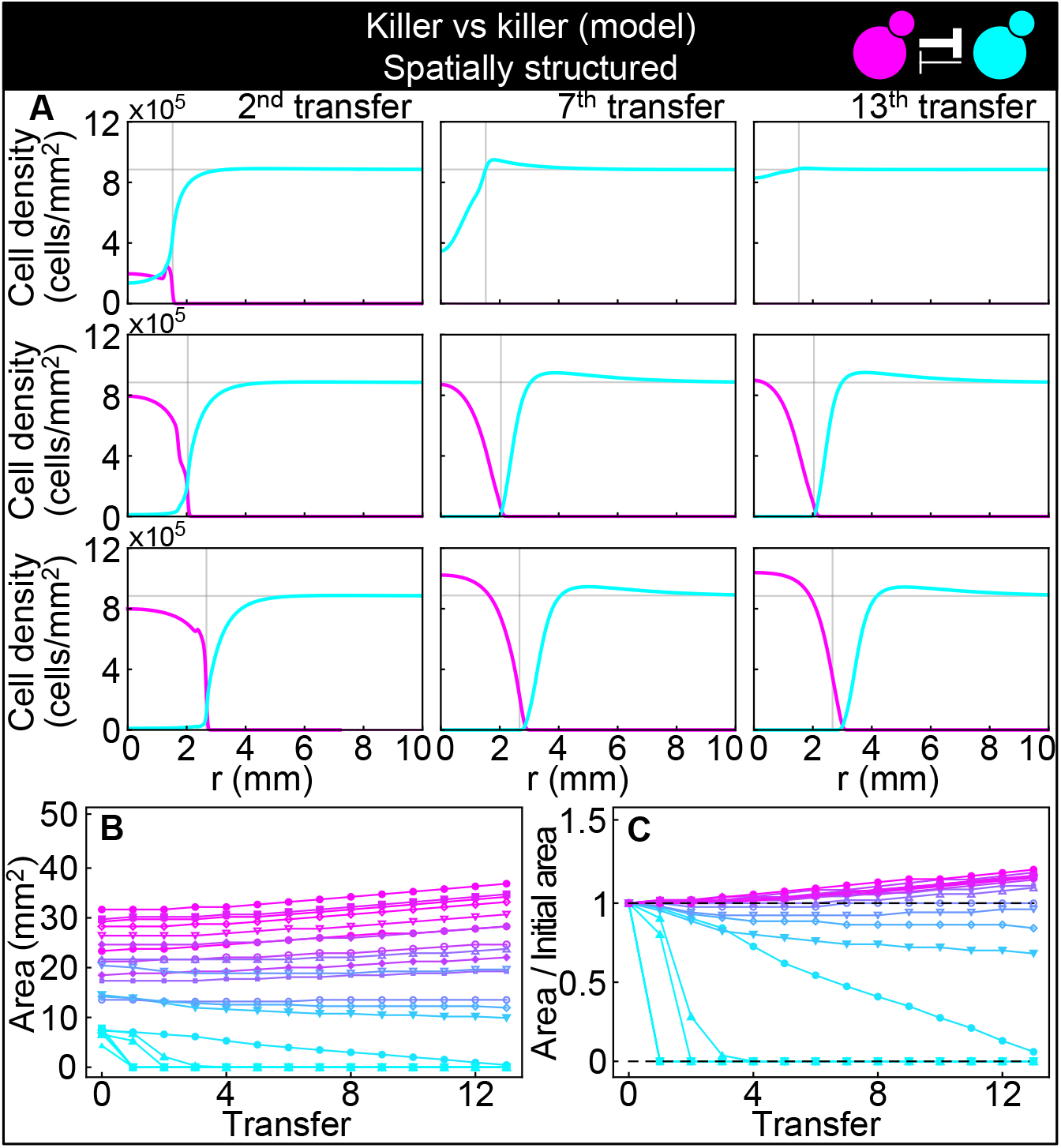
Numerical integrations of the spatial model (Equations 7–9), parametrized using the Experiments of Figure 2, and other values taken from the literature, can qualitatively reproduce the experimental dynamics of antagonistic competition between strains K1 and K2_b_ in spatially structured populations. **(A)** Simulated K1 inoculations smaller than the critical inoculum (first row) expanding on a landscape occupied by strain K2_b_ fail to establish and expand, whereas larger inoculations do (second and third row). The model can reproduce the formation of the halo, a region without cells at the interface between the two strains (second and third row). **(B)** Area covered by each simulated K1 population expanding on a landscape occupied by strain K2_b_ at the end of each 48-h growth period between simulated transfers, color coded from cyan to magenta according to the total K1 population of each replica at the end of the first growth period. **(C)** Same data as in (B), divided by the initial area to highlight relative changes. Compare (B) and (C) to Figure 5C & D.

## Discussion

We constructed an experimental system to study the dynamics of antagonism between yeast strains that produce and release two different toxins. We used this system to study the dynamics of antagonism in zero-dimensional well-mixed, two-dimensional well-mixed, and two-dimensional spatially structured populations. We derived mathematical models to describe each of these scenarios, parametrized the models using competition assays between killer and nonkiller strains, and showed that these models could predict the dynamics of antagonistic competition between toxin-producing strains. We verified the theoretical prediction that a critical inoculum size is required for a stronger antagonist to invade and displace a resident population of a weaker antagonist, in all of the spatial and non-spatial scenarios we tested.

The experiments in which we inoculated small populations of a stronger antagonist on a landscape occupied by a weaker one (Figure 5) revealed two unexpected features of antagonistic dynamics that had not been theoretically considered or predicted before. First, the expansion of the stronger antagonist was preceded by a contraction phase in which a halo devoid of cells formed between the two antagonists, both of which initially retreated. The existence of a halo had been observed before and used as a readout for killer activity by yeast geneticists (Tipper & Bostian, 1984), but its temporal dynamics had not been investigated from the point of view of population dynamics. Our investigation of increasingly realistic spatial models suggests that the diffusion of nutrients away from the boundary between two antagonistic strains plays an important role in the formation of the halo. Following its initial retreat, as the halo formed, the stronger antagonist started to expand, while the weaker antagonist kept retreating. Second, we found that occasionally new phenotypes emerged in the experiments after repeated transfers of the populations. Specifically, cells of the weaker, resident antagonist, resistant to the toxin produced by the stronger one, emerged in the population and formed fronts that re-invaded the population of the stronger antagonist. We believe that resistant cells were able to cross the nutrient-depleted region of the halo because, right after the populations were diluted by replica plating, resistant cells could grow and divide despite the presence of the invader’s strain toxin, and they could thus take up nutrients located in that region of space before those nutrients diffused away. We also observed a population of the stronger antagonist with reduced killing activity, showing that the strength of the antagonistic interaction varied throughout the experiments and raising the interesting question of how such a mutant succeeded in outcompeting its ancestor which made more toxin. A recent study found that a strain infected with the K1 killer virus first lost the ability to produce the toxin, and later lost immunity to the K1 toxin during an evolutionary experiment which lasted for 1,000 generations in well-mixed liquid cultures (Buskirk, Rojes, & Land, 2020). In that context, immunity to the toxin was lost once the killer toxin was not present in the medium anymore. In our experiments, the outlier population K1^o^ reduced its killing activity while the toxin produced by the resident strain was still present, suggesting that the evolutionary dynamics of microbial antagonism is quite rich and includes features that we do not fully understand.

Our results for yeast warfare may also have implications for interactions between two antagonistic strains carried on the surface of liquid substrates. When confined at air-liquid interfaces due to capillary forces, the metabolism of *S. cerevisiae* growing on a viscous liquid can produce density changes that generate fluid flows many times larger than their unperturbed colony expansion speed. That flow, in turn, can dramatically impact colony morphology and spatial population genetics (Atis, Weinstein, Murray, & Nelson, 2019). In these situations, an energetic cost associated with the interface between the area occupied by cells and the surrounding liquid (line tension) can play an important role, especially when a metabolically-induced flow beneath the colony leads to the extrusion of thin strands of the colony (a fingering instability). For such surface-borne communities, the combination of fluid flow and line tension can have a profound effect on genetic outcomes. Our antagonistic yeast strains could be used to ask what happens to range expansions on liquid or solid substrates when the density dependence of killing generates a form of line tension for each strain and antagonism generates halos of destruction between them.

There is growing interest in designing synthetic microbial consortia made of different microbial strains that interact to produce various tasks (Kong, Meldgin, Collins, & Lu, 2018). From the engineering perspective, these synthetic consortia may be valuable tools to manipulate microbiomes for environmental and health-related applications. For example, using antagonistic interactions has been proposed as a strategy for antimicrobial intervention (Gonzalez, Sabnis, Foster, & Mavridou, 2018). Synthetic consortia with predefined social interactions and some degree of control over the strength of such interactions (usually stepwise by using different promoters) have been designed using for example the prokaryotes *Lactococcus lactis* (Kong et al., 2018) and *Escherichia coli* (Ozgen et al., 2018), and commensalism and cooperation have been designed in *S. cerevisiae* (Muller et al., 2014), where the strength of the interaction has been controlled in a continuous way by varying the availability of shared resources in the extracellular medium. Our work provides a synthetic model of microbial antagonism in which the interaction strength of two antagonist strains can be controlled independently and continuously, via titratable induction of promoters that respond to different chemicals. There are other killer yeast viruses and corresponding toxins available (Liu et al., 2015; Magliani et al., 1997; Schmitt & Breinig, 2006), as well as inducible promoters that respond to chemicals other than copper and galactose (Lindstrom & Gottschling, 2009; Sangsoda, Cherest, & Surdin-Kerjan, 1985), leaving the possibility of expanding this system to more than two antagonist strains in the future. The ability to use engineered microbes to understand the fundamental features of antagonistic dynamics in simple spatial and non-spatial settings will help to model and design antagonistic interactions that could be exploited in synthetic microbial consortia, for example to ensure that a strain can invade a pre-existing consortium and persist therein.

Our results may be of interest beyond the spatial dynamics of antagonistic microbial populations. Laboratory experiments with yeast and bacteria growing on surfaces have increased our understanding of ecological and evolutionary dynamics in dense microbial populations and biofilms (Giometto et al., 2018; Hallatschek et al., 2007; Kayser et al., 2019; Kayser et al., 2018), but also have implications for other dense cellular populations such as tumors growing in three dimensions (Lamprecht et al., 2017; Lavrentovich & Nelson, 2015). Cancers made of heterogeneous clonal populations, in particular, display a rich set of interactions among clonal sub-populations including one-way antagonistic interactions in which one clone inhibits others (Marusyk & Polyak, 2010), a situation analogous to the one considered in the experiments of Figure 2. Of course, the interaction dynamics and geometry of heterogeneous clonal populations in cancer is much more complex than the experiments performed here. The spatial dynamics of antagonistic populations (Lavrentovich & Nelson, 2019), as well as those of gene drives (Tanaka et al., 2017), and of hybrid zones (Barton & Hewitt, 1985; Rouhani & Barton, 1987; Szymura & Barton, 1986) all share analogies with classical nucleation physics. For all these spatial processes, one can define a potential energy function (Figure 3C) with two minima corresponding to stable points in which one variant competitively excludes the other one, separated by a finite energy barrier that defines a critical inoculum size required to establish a successful, expanding invasion that drives the interacting system from one state to the other. In the case of gene drives, such an energy barrier may help preventing the accidental spread of these genetic constructs (Tanaka et al., 2017), which have the potential of damaging natural ecosystems.

Finally, we note that our mathematical models predict the behavior of idealized and unchanging populations, whereas organisms acquire mutations and selection on these mutations can produce results that violate simple, theoretical predictions. In our experiments, we saw the appearance of mutations that altered the outcome of competitions by either changing the rate of toxin production or the sensitivity to the toxin secreted by an antagonistic strain. In the natural world, the appearance and interactions of such mutants is likely to play a substantial role in the long-term behavior of antagonistic populations, especially in spatially structured populations where the descendants of the original mutant cell remain close to each other and can reap collective benefits from their altered behavior.

## Methods

Kev Resources Table.

| Key Resources Table |  |  |  |  |
| --- | --- | --- | --- | --- |
| Reagent type (species) or resource | Designation | Source or reference | Identifiers | Additional information |
| Strain, strain background (*S. cerevisiae*) | W303 | Murray lab | W1588 | The complete list of derived strains is available in Table 2 |
| Chemical compound, drug | Copper(II) sulfate pentahydrate | Sigma Aldrich | C8027 |  |
| Chemical compound, drug | D-(+)-Galactose | VWR | VWRV0637 |  |
| Chemical compound, drug | ethylene glycol-bis(β-aminoethyl ether)-N,N,N',N' | Sigma Aldrich | E34378 | |
|  | tetraacetic acid (EGTA) |  |  |  |
| Software, algorithm | Fiji (ImageJ) | PMID: 22743772 |  |  |
| Software, algorithm | FlowCytometryTools | https://eyurtsev.github.io/FlowCytometryTools/ |  | Version 0.5.0 |
| Software, algorithm | Mathematica | Wolfram | SCR_014448 | Version 12.1.0.0. Custom scripts available at https://github.com/andreagiometto/Giometto_Nelson_Murray_2020 |
| Software, algorithm | Matlab | Mathworks | RRID:SCR_001622 | Version R2018a. Custom scripts available at https://github.com/andreagiometto/Giometto_Nelson_Murray_2020 |
| Software, algorithm | Python | Python | RRID:SCR_008394 | Custom scripts available at https://github.com/andreagiometto/Giometto_Nelson_Murray_2020 |

### Genetic constructs

The killer toxin and fluorescent protein genes used in this study were cloned into integrative plasmids. The K1 killer toxin gene was PCR amplified from the plasmid YES2.1/V5-HIS-TOPO- K1 pptox (Breinig, Sendzik, Eisfeld, & Schmitt, 2006; Gier, Schmitt, & Breinig, 2017), which contains a DNA copy of the region of the dsRNA M1 virus genome encoding for the K1 toxin, and for the immunity of the host cell to the toxin, followed by the terminator *T_CYC1_*. We obtained YES2.1/V5-HIS-TOPO-K1 pptox from Manfred Schmitt and Frank Breinig. The primers used for the amplification were oAG24 and oAG25, which were designed to clone the *P_GAL1_* promoter in front of the K1 killer toxin gene via Gibson assembly, and to keep *T_CYC1_* at the end of the K1 killer toxin gene. The K2 killer toxin gene was amplified via PCR from a pUC-based plasmid containing a DNA copy of the region of the dsRNA M2 virus genome that encodes for the K2 toxin (Servienė, Čepononytė, Lebionka, & Melvydas, 2007), and for the immunity of the host cell to the toxin. We obtained this pUC-based plasmid from Elena Servienė. The primers used were oAG26 and oAG29, which were chosen to clone the *P_CUP1_* promoter in front of the K2 killer toxin gene and the terminator *Tcyci* after it via Gibson assembly. The terminator *T_CYC1_* was amplified via PCR from YES2.1/V5- HIS-TOPO-K1 pptox using primers oAG25 and oAG31. The promoter *P_GAL1_* was amplified via PCR with primers oAG22 and oAG23 from a *pFA6a-kanMX6-PGALi* plasmid (Longtine et al., 1998; Wach, Brachat, Pohlmann, & Philippsen, 1994). The promoter *P_CUP1_* was amplified via PCR with primers oAG27 and oAG28 from a *pFA6a-Pcupi-UBI-DHFR* plasmid. The segments containing *P_GAL1_* and *K1-T_CYC1_* were assembled via Gibson assembly (plasmid pAG11), using as backbone a *pFA6a-prACT1-ymCherry-KanMX6* plasmid (pAG3) linearized with the restriction enzymes *EcoRI* and *EcoRV.* The segments containing *Pcupi, K2* and *T_CYC1_* were assembled via Gibson assembly (plasmid pAG14), using as backbone a *pFA6a-prACT1-ymCitrine-KanMX6* plasmid (pAG5) linearized with the restriction enzymes *EcoRI* and *EcoRV.* These plasmids were linearized at the *T_CYC1_* locus using the restriction enzyme *PpuMI,* and their integration at the *T_CYC1_* locus was verified using colony PCR using primers oAG5/oAG46 and oAG44/oAG45 (strain K1), and oAG8/oAG46 and oAG44/oAG45 (strains K2 and K2_b_).

### Strains

The strains used here were derivatives of the *S. cerevisiae* strain yJHK234 derived from the W303 genetic background. This strain was constructed as described in (Ingolia, 2006; Ingolia & Murray, 2007) such that expression from *Pgal1* occurs in a titratable, unimodal way in response to changes in the extracellular concentration of galactose, because galactose has been turned into a gratuitous, non-metabolizable inducer by deleting the genes *GAL1* and *GAL10* from its genome, and the bistability in galactose induction has been removed by placing the *GAL3* gene under the constitutive promoter *PACT1* (Ingolia, 2006). By competing yJHK234 against reference K1 (strain F166), K2 (strain EX73), K28 and nonkiller K^−^ strains obtained from Manuel Ramírez (Maqueda, Zamora, Alvarez, & Ramirez, 2012), we found that yJHK234 has a K1^+^ phenotype, being resistant to the toxin produced by the reference K1 strain, sensitive to the toxins produced by the reference K2 and K28 strains, and capable of killing the reference nonkiller K^−^ strain. For our purposes, we needed an ancestor strain cured of the M1 virus, which could serve as an ancestor for both the killer, toxin-producing strains and for the sensitive, nonkiller ones. To obtain clones cured of the M1 virus, we spread about 100 cells on YPD agar plates with pH 4.5 and 0.001% (w/v) methylene blue (an indicator for cell death), incubated at 25°C for 24 h and isolated blue colonies (clones cured of the virus being killed by surrounding non-cured colonies). We tested that the isolated clone yAG74 was sensitive to toxins secreted by both the K1 and K2 reference strains. To prevent catabolite repression, which would prevent expression of the K1 toxin from *P_GAL1_* in the presence of glucose, we deleted the hexokinase 2 gene, *HXK2,* (Raamsdonk et al., 2001) from yAG74 (leading to yAG75) via transformation of the *HphMX4* marker from a *pFA6-HphMX4* plasmid with 5’ and 3’ flanking sequences of *HXK2,* using primers oAG13 and oAG14. We checked the deletion using colony PCR with primers oAG15/oAG17 and oAG15/oAG39. The K1 killer strain (yAG94) was obtained by transforming the linearized plasmid pAG11 into strain yAG75. The K2 (yAG83) and K2_b_ (yAG82) killer strains were obtained by transforming the linearized plasmid pAG14 into strain yAG75. The transformant clones yAG94 and yAG83 were chosen for the experiments because of the bright fluorescence signal of the transformed cells observed at the flow cytometer and at the stereomicroscope compared to other transformants, which suggests multiple integrations of the plasmid into the genome. Conversely, yAG82 was selected because of the weaker fluorescent signal of the transformed cells compared to yAG83, suggesting a single integration of the plasmid. The sensitive, nonkiller strains S1 (yAG96) and S2 (yAG99) were obtained by transforming strain yAG75 with the linearized plasmids pAG3 and pAG5 digested with the restriction enzyme *AgeI* (which cleaves DNA in *P_ACT1_*), respectively.

### Media and growth conditions

All experiments were performed using YPD buffered at pH 4.5 and supplemented with adenine, tryptophan and ethylene glycol-bis(β-aminoethyl ether)-N,N,N’,N’- tetraacetic acid (EGTA), a chelating agent that we used to reduce the baseline expression of *P_CUP1_*. The medium was prepared by mixing 990 mL of Millipore-purified water, 20 g of BD Bacto Peptone, 10 g of BD Bacto Yeast Extract, 10 mL of a 1% (w/v) solution of adenine and tryptophan, 1.04 g of NaOH and 9.51 g of EGTA. Then, 2 g of NaOH were added to bring the EGTA into solution. Then, we added 11.2 g of succinic acid and brought the pH to 4.5 by adding approximately further 2 g of NaOH. The solution was then filter-sterilized. Agarose medium was prepared following the same procedure but using 590 mL of water instead of 990 mL. Separately, 400 mL of Millipore water were mixed with 20 g of BD Bacto Agar and microwaved for 2 min. The two solutions were then combined and used to fill Petri dishes. Solutions of copper(II) sulfate and of galactose at different concentrations were added to the media in different volumes according to the desired final concentration of the two inducers.

### Competition assays

Competition assays in liquid media were performed as follows. Strains were plated from the glycerol stock four days prior to the start of the experiment and grown for two days at 30°C in YPD plates. One day before the start of the competition assays, overnight cultures were started by transferring cells from the plates to a tube containing 2 mL YPD buffered at pH 4.5, which was placed in a rotating roller drum at 30°C. At the start of the competition assays, 200 uL from the overnight cultures were centrifuged, the supernatant was removed, and cells were then resuspended in 2 mL autoclaved water. The centrifugation and resuspension were repeated twice to dilute away any toxins produced overnight. For killer-vs-nonkiller competition assays, the strains K1, K2 and K2_b_ were mixed with strains S2, S1 and S1, respectively (so that the two strains expressed different fluorescent proteins). Cell suspensions containing different strains were then mixed at the desired relative frequencies, and 40 uL of these were then diluted in 10 mL YPD buffered at pH 4.5 with EGTA. Each replica in the competition assays consisted of 500 uL of this solution placed in deep-welled (capacity 2 ml/well), 96-well, round-bottomed plates, taped to a roller drum rotating at a frequency of 1 rotation per second and placed in a room kept at 25°C. Technical replicates were assigned to random positions on the 96-well plate, irrespective of the treatment they belonged to. At regular time intervals, small samples (≤10 uL) were taken from each well, diluted in 50 mM TRIS-HCl, pH 7.8 and the relative frequencies of the two strains was measured by flow cytometry. Flow cytometry data was performed using the Python package FlowCytometryTools and custom Python and Mathematica scripts. Occasionally, during measurement with the flow cytometer, some wells were not measured due to the aspiration of bubbles by the robotic liquid handler that automatically measured the 96 well plates. Due to the temporal sensitivity of the assay, relative frequency data from those replicates could not be recovered, and thus we excluded those technical replicates from the analysis. Competition assays with the unusual cells sampled from the experiments of Figure 5 (assays of Figure 6) were performed using the same protocol as the competition described above, starting from glycerol stocks of the cells sampled from the experimental populations of Figure 5.

### Spatial experiments

The experiments with spatially well-mixed populations on surfaces shown in Figure 4 were performed as follows. 28 mL of molten YPD agar with pH 4.5 and EGTA *(Media and growth conditions)* were added to 100 mm diameter Petri dishes two days before the start of the experiment, along with appropriate amounts of a 5 mM solution of copper(II) sulfate or a 50 mM solution of galactose to reach the desired target concentration of inducer on the plates. The day before the experiment we inoculated overnight cultures of strains K1, K2, S1 and S2 in 2 mL YPD culture tubes with pH 4.5 and 25 mM EGTA and grew them at 30°C on a rotating roller drum. At the start of the experiment, we centrifuged and resuspended 200 uL of the overnight cultures in 2 mL autoclaved Millipore-purified water, repeating the centrifugation and resuspension twice to remove toxins from the overnight cultures. We mixed strains K1 and K2 with K1 frequencies *f = 1/2*,3/5, *7/10, 4/5* and *9/10* for the treatment with 12.5 uM copper, with K1 frequencies *f = 1/5,7/20, 1/2, 13/20* and *4/5* for the treatments with 250 uM, 270 uM and 300 uM galactose, and with K1 frequencies *f = 1/10*,1/5, *3/10, 2/5* and *1/2* for the treatment with 350 uM galactose. For the control experiments (insets in Figure 3B), we mixed strains S1 and S2 with S1 frequencies *f = 1/10*, *3/10, 1/2, 7/10* and *9/10* for both treatments (12.5 uM copper and 350 uM galactose). 5 uL droplets of the mixed solutions were inoculated at different, random locations on a regular lattice on the surface of the agar. Plates were placed inside a plastic box with an open water Schott flask to provide humidity, in a room set at 25°C. When depositing a droplet from an overnight culture on a plate, a large fraction of the cells in the droplet end up at distributed at the outer boundary of the droplet due to the coffee-stain effect (Deegan et al., 1997). In this experiment, we imaged the interior of the droplets deposited on the agar surface, far from the coffee-stain ring. The experiments with spatially structured populations on surfaces shown in Figure 5 and its figure supplements were performed similarly but imaging the entire droplets. At the start of those experiments, we centrifuged and resuspended 200 uL of the overnight cultures K1, K2_b_, S1 and S2 in 200 uL autoclaved Millipore water, repeating the centrifugation and resuspension twice to remove toxins from the overnight cultures. For the experiments of Figure 5, 200 uL of the resuspended overnight K2_b_ culture were spread on the surface of the agar using an inoculating loop. Then, using a micropipette, we deposited droplets of the resuspended overnight K1 culture on top of the K2_b_ lawn, at random locations on a regular lattice, well separated from each other. We deposited droplets of 6 different volumes (0.5 uL to 3 uL with 0.5 uL increments), with 7 replicates per volume, across multiple plates. Similarly for the experiments shown in Figure 5 - supplement 2, 200 uL of the resuspended overnight S2 culture were spread on the surface of the agar using an inoculating loop, and droplets of volumes 0.5 uL, 2 uL and 3 uL of the S1 culture were deposited on top of the S2 lawn, with 7 replicates per volume, at random locations on a regular lattice. We imaged each population at the end of the each 48-h growth period, immediately before its transfer to the next plate, using a fluorescence stereomicroscope, always at the same magnification and with the same exposure time. The areas of the invading populations shown in Figure 5C-D and Figure 5 - supplement 2B-C were measured via image analysis using custom Fiji scripts and Mathematica notebooks.

### The frequency model

The starting point for the derivation of the frequency model (Equation 1) is the following set of equations for the densities of two toxin-producing strains (*n*_1_ and *n*_2_) and the concentrations of the two toxins they produce (*c*_1_ and *c*_2_, respectively):

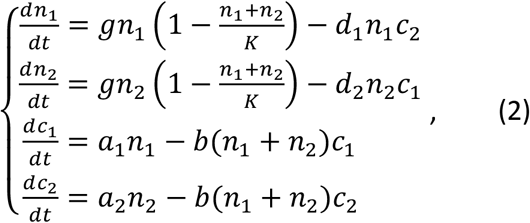

where *n_i_* is the cell density of strain *i, g* is the growth rate of the two strains in isolation (assumed to be identical for the two strains as they differ solely for the integrated genetic construct), *c_i_* is the concentration of toxin *i, d_i_*s are death rates per concentration of *c_j_* (*j* ≠ *i*), and *a_i_*s and *h_i_*s are toxin production and toxin attachment rates (the two toxins bind to the same receptor on the cell wall of both sensitive and producer strains, and thus the term *n*_1_ + *n*_2_). Equations (2) assume that the growth of each strain can be described by a logistic growth term in which the carrying capacity is shared by the two strains. Toxin-induced cell death is introduced via a term proportional to the product between a strain’s density and the concentration of the toxin produced by the other strain, and toxin production is assumed to be proportional to the density of the strain that produces it. Upon assuming that *c*_1_ and *c*_2_ are “fast variables” (in a sense specified below) compared to *n*_1_ and *n*_2_, we set their temporal derivatives to zero to find the quasistatic approximations *c*_1_ = *a*_1_*n*_1_/[*b*(*n*_1_ + *n*_2_)] and *c*_2_ = *a*_2_*n*_2_/[*b*(*n*_1_ + *n*_2_)]. After substituting these approximations in the first two lines of Equations (2) and computing the temporal derivative for the fraction of strain 1 in the population, *f* = *n*_1_/(*n*_1_ + *n*_2_), we find:

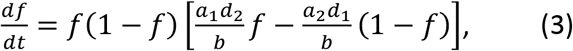

which is in fact Equation (1) with the interaction coefficients *r*_1_ = *a*_1_*d*_2_/*b* and *r*_2_ = *a*_2_*d*_1_/*b*. We can rewrite Equation (3) as:

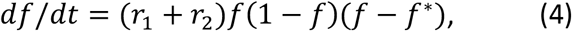

with *f* * = *r*_2_/(*r*_1_ + *r*_2_). Equation (4) reveals that the dynamics of *f* evolves with the characteristic time scale *τ*. = 1/(*r*_1_ + *r*_2_). Once we indicate with *c*_1_ and *c*_2_ the quasi fixed points *c*_1_ = (*a*_1_/*b*)*f* and *c*_2_ = (*a*_2_/*b*)(1 – *f*), and letting *c*_1_(*t*) = *c*_1_ + *δc*_1_(*t*) and *c*_2_(*t*) = *c*_2_ + *δc*_2_(*t*), we can rewrite the last two lines of Equations (2) as:

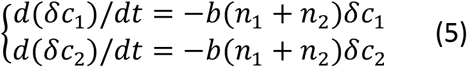

which show that the toxin concentrations evolve with the characteristic time scale *τ_tox_* = 1/(*bn*), with *n*_1_ = *n*_2_ + *n_2_*. Thus, for the quasistatic approximation to hold we require that *τ_tox_* ≪ *τ_f_*., i.e. that *r*_1_ + *r*_2_ ≪ *bn*. A best fit of Equation (2) to the data on the competition between toxin-producing and sensitive, nonkiller strains allow us to check if this condition is met. For example, for the competition of K1 versus K2 with 360 uM galactose and 0 uM copper we have *r*_1_ = 0.13 h^−1^, *r*_2_ = 0.04 h^−1^ and *b* = 1.2 · 10^−8^ h^−1^ (cell/mL)^−1^ (see next section for its estimate), and thus the condition is satisfied for *n* ≫ (*r*_1_ + *r*_2_)/*b* ≈ 10^7^ cells/mL, which we can compare to the carrying capacity *K* = 2.3 · 10^8^ cells/mL. With a starting density of about 3 · 10^4^ cell/mL and a growth rate *g* = 0.35 h^−1^, it takes about 17 h for the condition *n* » (*r*_1_ + *r*_2_)/*b* to be met, and thus the frequency model is only strictly appropriate for describing the last few hours of the dynamics of our competition assays. This might explain why the frequency model tends to slightly overestimate the increase in frequency at *t* = 14 h of the strongest antagonist strain in our competition assays, with respect to the data (Figures 2A and 3A). Nonetheless, the frequency model seems to do a good job in predicting the antagonistic dynamics between toxin-producing strains, possibly because the early phases of the dynamics are dominated by the exponential growth of the two strains and the toxins don’t affect the dynamics too much at this stage. Strictly speaking, when the condition *n* ≫ (*r*_1_ + *r*_2_)/*b* is not met, the full model in Equations (2) would be better suited to describe the data.

### Spatial models

Our starting point for the investigation of antagonistic population dynamics in spatially structured populations was the following spatial generalization of Equations (2):

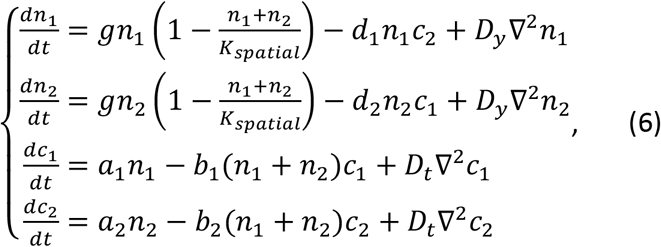

where the constants *D_y_* and *D_t_* are the diffusion rates of the yeast cells and their toxins (the two toxins K1 and K2 have similar sizes, so we assumed that they have identical diffusion rates), and *K_spatial_* is the spatial carrying capacity (with units of cells/cm^2^), which is related to the well-mixed carrying capacity via the relationship *K_spatial_ = Kh*, where *h* is the height of the medium in the agar plate. We used a Markov-Chain-Monte-Carlo (MCMC) algorithm (Vrugt et al., 2009) to fit the non-spatial version of this model (Equations 2) to the data from the competition assays between toxin-producing and nonkiller strains. To reduce the number of free parameters, we assumed *b*_1_ = *b*_2_ and we partially nondimensionalized the equations through appropriate rescaling of the variables and parameters (section *Parameter fitting).* We found that the non-spatial version of this model could indeed fit the antagonistic dynamics between toxin-producing and nonkiller strains, and that it could predict the dynamics of antagonistic competition between the two toxin-producing strains K1 and K2 (and K1 versus K2_b_) in well-mixed media (Figure 3 - figure supplement 1). However, when we numerically integrated the model in an attempt to reproduce the dynamics that we had observed in the spatially structured experiments, Equations (6) failed to reproduce the halo between the two toxin-producing strains (Figure 7 - figure supplement 1A), at least within a reasonable range for the various parameters of the model. The failure of such model to reproduce the halo can be explained as follows. The logistic growth term in Equations 6 assumes that every cm^2^ on the agar can support *K_spatial_* cells. In such a model, nutrients located in a given region of space cannot diffuse to nearby regions and thus can only support the growth of cells locally. With toxin production rates representative of our experiments, the toxin produced by an antagonist strain is not sufficient to completely halt the growth of the other antagonist, as shown by the fact that the absolute number of cells of both antagonists grew in all our well-mixed competition experiments, even if the relative frequency of one of the strains declined with time. In the model with logistic growth, the two populations are thus able to grow at the interface between the two antagonist strains, even if at a slower pace compared to other regions of space, eventually almost completely filling the halo region with cells (Figure 7 - figure supplement 1A). When nutrients can diffuse, however, nutrients move to other regions of space before cells at the interface between the two antagonists are able to grow, leading to the depletion region that we referred to as the ‘halo’. Variants of Equations (6) that accounted for a more realistic cell diffusion term (see discussion below Equations 7), for the fact that the toxin production rate is likely dependent on growth rate and for the fact that the toxins can diffuse into the agar below the surface on which cells are located also failed to reproduce the halo using biologically realistic parameters (Figure 7 - figure supplement 1B-D). These results motivated us to gradually increase the complexity and realism of the model to identify the processes that are responsible for the dynamics observed in the experiments. In the most realistic model we investigated, the equations for the two strains, inhabiting the two-dimensional surface at the top of the agar (at the height coordinate *z* = *h*), read:

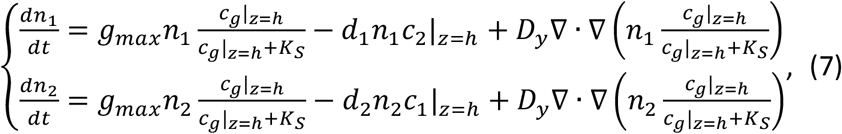

where *c_g_*|_*z=h*_ is the glucose concentration (*c_g_*) at the agar surface (|_*z=h*_), *K_s_* is Monod’s half-saturation constant for the growth rate of *S. cerevisiae* on glucose, *g_max_* is the maximum growth rate of the two strains (attained in the limit of infinite glucose concentration), and *c _i_*|_*z=h*_ is the concentration of toxin *i* at the agar surface. Compared to Equation 6, the diffusion term has been modified to reflect the fact that in our experiments the predominant contribution to the local diffusion of cells is not due to Brownian motion, but rather to the growth dynamics of mother cells giving rise to daughter cells in their immediate surroundings, and to the excluded volume forces that cells exert on each other while growing and dividing (Giometto et al., 2018; Kayser et al., 2019). To model this phenomenon, we took the local flux of cells to be proportional to the local growth rate (having a standard diffusion term in Equations (7) does not alter the results significantly, but it seems to us more realistic to have a growth-rate dependent diffusion term in the model). The dynamics of *c*_1_, *c*_2_ and *c_g_* are now governed by the diffusion equation within the agar (0 ≤ *z < h*):

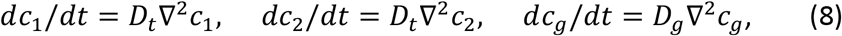

where *D_g_* is the diffusion coefficient of glucose, complemented by the following fluxes at the agar surface *z = h*:

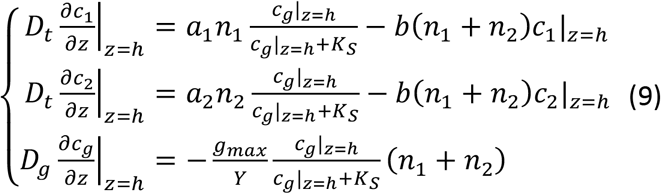

Here *Y* is the cellular yield (cells/g glucose), and we impose no-flux boundary conditions on all other surfaces (i.e., where the agar is in contact with the Petri dish plastic). We parametrized the non-spatial version of this model using an MCMC algorithm (Vrugt et al., 2009) and the data from the competition assays between the killer strains and the nonkiller strains S1 and S2. We found that the non-spatial version of this model could fit the antagonistic dynamics between toxin-producing and nonkiller strains, and that it could also predict the dynamics of antagonistic competition between the two toxin-producing strains K1 and K2 (and K1 vs K2_b_) in well-mixed media.

### Parameter fitting

The parameters of the frequency model were obtained by least-squares fitting of the equation:

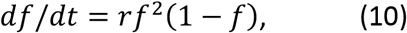

where *f* is the frequency of the toxin-producer strain, to the data on the competition between toxin-producing and sensitive, nonkiller strains shown in Figure 2. Note that Equation (10) is a special case of Equation (1) and describes a toxin-producer strain competing against a nonkiller one. The best-fit parameters obtained via least-squares fitting are reported in Table 3.

The parameters of Equation 2 were fit to the cell density data from the competition assays between toxin-producing and sensitive, nonkiller strains using MCMC (Vrugt et al., 2009). For a toxin-producing (K) strain competing versus a sensitive, nonkiller strain (S), Equation (2) reads:

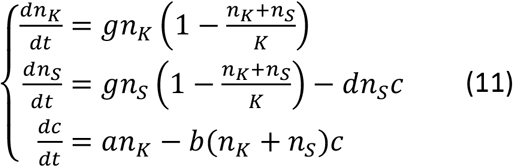

where *c* is the toxin concentration. Not all parameters in Equation (11) are identifiable, thus we rescaled the toxin concentration as *c*’ = *dc* and the toxin production rate as *a*’ = *da*, which is formally equivalent to setting *d* = 1 in Equation (11), although the measurement units are affected as the rescaled toxin concentration has dimensions of 1/time. Apostrophes are dropped in the following for clarity. Note that the toxin production rate depends on the inducer concentrations, and thus we have a value of *a* for each inducer concentration used. The initial condition for the relative frequencies of the two strains in the numerical integrations of Equation (11) was assumed to be equal to the relative frequencies at the first measurement time point, i.e. we assumed that the toxin did not alter the relative frequency of the two strains from inoculation to the first measurement time point. The assumption is justified because both the total cell density and the toxin concentration (*c* = 0 at *t* = 0) are low in the first phases of the dynamics. The initial condition for the total cell density *n*_0_, for each choice of *g* and *K* in the Markov Chain, was set to 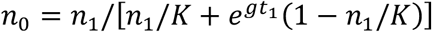, which is the solution of the logistic equation 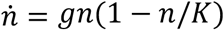 for *n*_0_, with *n*(*t*_1_) = *n*_1_ where *t_r_* = 14 h is the first measurement time point and *n*_1_ was taken from the data. All data were fit simultaneously, because the parameters *g, K* and *b* appear for all inducer concentration treatments. The best fit parameters are given in Table 4. Note that when comparing the predictions of this model to the data on the antagonistic competition between two killer strains, some of the parameters appear for more than one competition assay. For example, the parameter 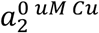 appears in all the competitions between K1 and K2 in which we added only galactose to the medium, and in the competition assay without any inducer.

**Table 4.**
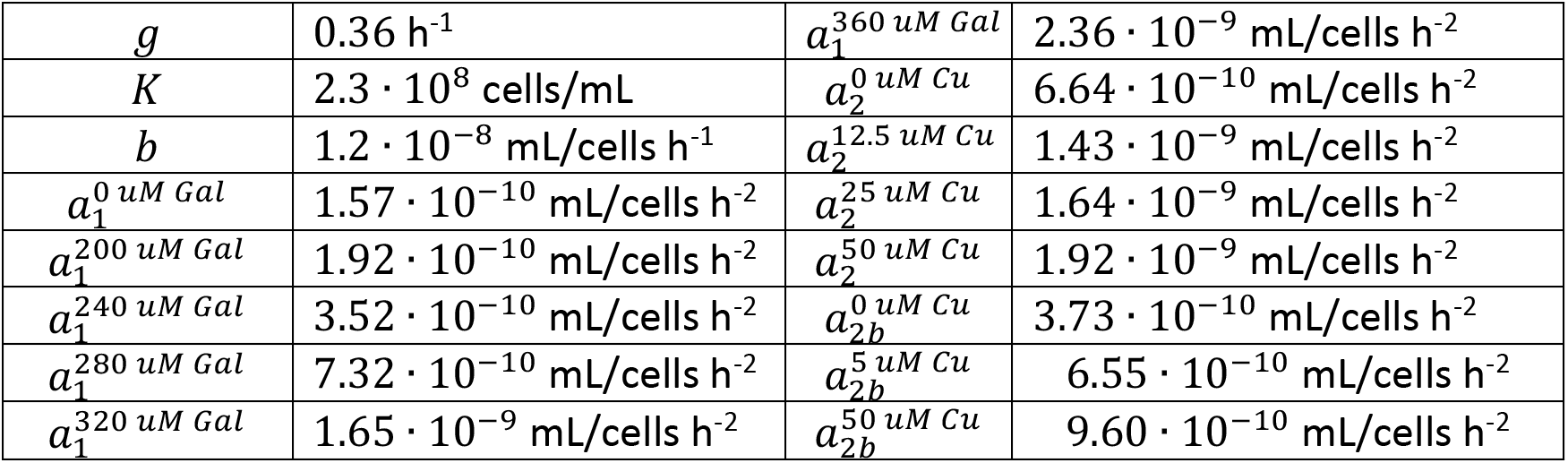
Best-fit estimates for the parameters of the full model with logistic growth (Equation 11) fitted to the data from competition assays between toxin-producing strains and sensitive, nonkiller ones, via MCMC. Concentrations of the inducers galactose (Gal) and copper (Cu) are indicated as superscripts. The parameters *a*_1_ and *a*_2_ correspond to the rescaled parameter *a*’ = *da* of Equation (11) for the strains K1 and K2, respectively.

|  |  |  |  |
| --- | --- | --- | --- |
| $g$ | $0.36 \text{ h}^{-1}$ | $a_1^{360 \text{ uM Gal}}$ | $2.36 \cdot 10^{-9} \text{ mL/cells h}^{-2}$ |
| $K$ | $2.3 \cdot 10^8 \text{ cells/mL}$ | $a_2^{0 \text{ uM Cu}}$ | $6.64 \cdot 10^{-10} \text{ mL/cells h}^{-2}$ |
| $b$ | $1.2 \cdot 10^{-8} \text{ mL/cells h}^{-1}$ | $a_2^{12.5 \text{ uM Cu}}$ | $1.43 \cdot 10^{-9} \text{ mL/cells h}^{-2}$ |
| $a_1^{0 \text{ uM Gal}}$ | $1.57 \cdot 10^{-10} \text{ mL/cells h}^{-2}$ | $a_2^{25 \text{ uM Cu}}$ | $1.64 \cdot 10^{-9} \text{ mL/cells h}^{-2}$ |
| $a_1^{200 \text{ uM Gal}}$ | $1.92 \cdot 10^{-10} \text{ mL/cells h}^{-2}$ | $a_2^{50 \text{ uM Cu}}$ | $1.92 \cdot 10^{-9} \text{ mL/cells h}^{-2}$ |
| $a_1^{240 \text{ uM Gal}}$ | $3.52 \cdot 10^{-10} \text{ mL/cells h}^{-2}$ | $a_{2b}^{0 \text{ uM Cu}}$ | $3.73 \cdot 10^{-10} \text{ mL/cells h}^{-2}$ |
| $a_1^{280 \text{ uM Gal}}$ | $7.32 \cdot 10^{-10} \text{ mL/cells h}^{-2}$ | $a_{2b}^{5 \text{ uM Cu}}$ | $6.55 \cdot 10^{-10} \text{ mL/cells h}^{-2}$ |
| $a_1^{320 \text{ uM Gal}}$ | $1.65 \cdot 10^{-9} \text{ mL/cells h}^{-2}$ | $a_{2b}^{50 \text{ uM Cu}}$ | $9.60 \cdot 10^{-10} \text{ mL/cells h}^{-2}$ |

The parameters of Equations (7) and (9) were estimated by fitting the following model with Monod growth dynamics (Monod, 1949) to the data from competition assays between toxin-producing and sensitive, nonkiller strains:

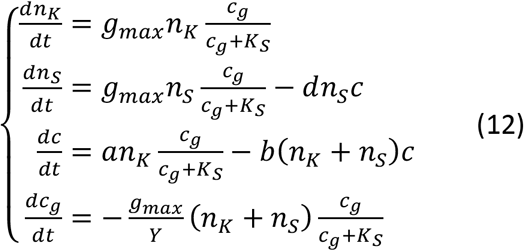

The toxin concentration and toxin production rate were rescaled as described previously. The initial concentration of glucose was set to the experimental value *c*_*g*0_ = 0.02 g/mL and the value for the Monod constant *K_s_* = 2 · 10^−5^ g glucose/mL (0.11 mM) was taken from the literature (Postma, Kuiper, Tomasouw, Scheffers, & Vandijken, 1989). The value of *K_s_* does not impact the results significantly within a biologically reasonable range, given that it only affects the later phases of the dynamics when glucose is depleted. The initial condition for the relative frequencies of the two strains in the numerical integrations of Equation (12) was assumed to be equal to the relative frequencies at the first measurement time point. The initial condition for the total cell density *n*_0_, for each choice of the other parameters in the Markov Chain, was set equal to the solution of the growth equation without antagonistic interactions 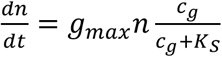 with 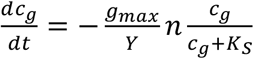 for *n*_0_ (here, *n = n_K_* + *n_s_*), which is also the solution of 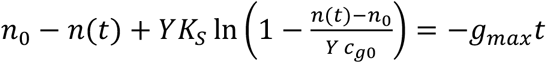 for *n*_0_ (we used the fact that in this simplified model *n* – *n*_0_ = *Y*(*c*_*g*0_ – *c*)), which we computed numerically using *t* = *t*_1_ = 14 h and *n*(*t*) = *n*_1_ taken from the data (first measurement time point). All data were fit simultaneously using MCMC, because the parameters *g_max_*, *Y* and *b* appear for all inducer concentration treatments. The best fit parameters are given in Table 5. Also for this model, when comparing the prediction of the model to the data on the antagonistic competition between two killer strains, some of the *a* parameters appear for more than one competition assay.

**Table 5.** Best-fit estimates for the parameters of the model with Monod growth dynamics (Equation 12) fit to the data from competition assays between toxin-producing strains and sensitive, nonkiller ones, via MCMC. Concentrations of the inducers galactose (Gal) and copper (Cu) are indicated as superscripts. The parameters *a*_1_, *a*_2_ and *a*_2*b*_ correspond to the rescaled parameter *a*’ = *da* in Equation 12 for the strains K1, K2 and K2_b_, respectively.

|  |  |  |  |
| --- | --- | --- | --- |
| $g_{max}$ | 0.31 1/h | $a_1^{360 \text{ uM Gal}}$ | $1.76 \cdot 10^{-9}$ mL/cells h<sup>-2</sup> |
| $Y$ | $1.3 \cdot 10^{10}$ cells/(g glucose) | $a_2^{0 \text{ uM Cu}}$ | $5.09 \cdot 10^{-10}$ mL/cells h<sup>-2</sup> |
| $b$ | $3.2 \cdot 10^{-9}$ mL/cells h<sup>-1</sup> | $a_2^{12.5 \text{ uM Cu}}$ | $1.04 \cdot 10^{-9}$ mL/cells h<sup>-2</sup> |
| $a_1^{0 \text{ uM Gal}}$ | Set to 0 mL/cells h<sup>-2</sup> | $a_2^{25 \text{ uM Cu}}$ | $1.24 \cdot 10^{-9}$ mL/cells h<sup>-2</sup> |
| $a_1^{200 \text{ uM Gal}}$ | $1.65 \cdot 10^{-10}$ mL/cells h<sup>-2</sup> | $a_2^{50 \text{ uM Cu}}$ | $1.43 \cdot 10^{-9}$ mL/cells h<sup>-2</sup> |
| $a_1^{240 \text{ uM Gal}}$ | $2.45 \cdot 10^{-10}$ mL/cells h<sup>-2</sup> | $a_{2b}^{0 \text{ uM Cu}}$ | $3.47 \cdot 10^{-10}$ mL/cells h<sup>-2</sup> |
| $a_1^{280 \text{ uM Gal}}$ | $6.11 \cdot 10^{-10}$ mL/cells h<sup>-2</sup> | $a_{2b}^{5 \text{ uM Cu}}$ | $5.52 \cdot 10^{-10}$ mL/cells h<sup>-2</sup> |
| $a_1^{320 \text{ uM Gal}}$ | $1.18 \cdot 10^{-9}$ mL/cells h<sup>-2</sup> | $a_{2b}^{50 \text{ uM Cu}}$ | $7.57 \cdot 10^{-10}$ mL/cells h<sup>-2</sup> |

Confidence intervals for the model predictions in Fig. 3 were computed by plotting the model predictions using the interaction coefficients *r*_1_ ± *σ*_1_ and *r*_2_ ± *σ*_2_, where (*r*_1_ and *r*_2_ are the interaction coefficients best-fit estimates and *σ*_1_ and *σ*_2_ are the standard deviations reported in Table 3).

The diffusion coefficient of glucose in Equations (6–9) was taken from the literature, whereas the toxin diffusion coefficients were estimated based on experimental values for proteins of similar sizes (Magliani et al., 1997) diffusing in agar gels (Pluen, Netti, Jain, & Berk, 1999). Their values are reported in Table 6. The yeast diffusion coefficient was estimated as follows. The local movement of cells in our system is not predominantly due to Brownian motion, but rather to the forces that dividing cells exert on each other during growth. Thus, we estimated the yeast diffusion coefficient as *D_y_* = *d*^2^*g_max_*, where the typical yeast cell diameter *d* ≈ 10 um (estimated by measuring the mean cell diameter in liquid cultures of K1 and K2) and *g_max_* appear for dimensional reasons. Because cells are not locally in isolation and multiple cells are typically dividing and pushing each other at the same time, the effective value of *D_y_* may be larger in the experiments. Other factors that can contribute to increasing the effective diffusion coefficient are the collisions of local clusters of cells that are transferred from old to new plates with by replica plating, and the replica plating itself might also contribute given that the agar surface is pressed onto a microfiber cloth twice to transfer cells to a fresh plate. Increasing the value of *D_y_* in the simulations increases the critical inoculum size. For example, with *D_y_* = 3 ° 10^−6^ cm^2^/h (10 times larger than the value used here), the critical inoculum size is about 20 mm^2^. In Figure 7 - figure supplement 3 we used numerical simulations to investigate the dependence of the critical inoculum size on the toxin production rate of the invader and on the toxin diffusion rate in the model. We found that the critical inoculum size in our model is proportional to 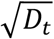, as shown by the fact that simulation data collapse onto a single curve when dividing the critical radius by this factor (Figure 7 - figure supplement 3B-C). Additionally, simulation data suggests that for large *a*_1_ – *a*_2_, the critical radius scales as 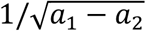 (Figure 7 - figure supplement 3C). In Figure 7 - figure supplement 4, we also show that the width of the halo varies with the toxin production rates of the two strains and with the diffusion coefficients of glucose and of the toxins.

**Table 6.** Values used in the numerical integrations of Equations (6–9) for the diffusion coefficients and Monod’s constant for growth of *S. cerevisiae* on glucose. The value of *K_s_* was taken from (Postma et al., 1989). As an estimate for the K1 and K2 toxins diffusion coefficient, we took a typical value for proteins of size similar to the K1 and K2 toxins (Magliani et al., 1997) diffusing in agar gels at 25°C (Pluen et al., 1999). The yeast diffusion coefficient *D_y_* was estimated as discussed in the text. The value for *D_g_* was taken from (Longsworth, 1955).

| $D_g$ | $D_t$ | $D_y$ | $K_S$ |
| --- | --- | --- | --- |
| 0.024 $\text{cm}^2/\text{h}$ | $3 \cdot 10^{-3}$ $\text{cm}^2/\text{h}$ | $3 \cdot 10^{-7}$ $\text{cm}^2/\text{h}$ | $2 \cdot 10^{-5}$ g glucose/mL |

### Numerical integration of the spatially structured model

We numerically integrated Equations (7) in polar coordinates and Equations (8–9) in cylindrical coordinates, using the forward Euler method and assuming symmetry in the azimuthal coordinate. The integration steps in the radial, altitudinal and temporal directions were set to *dr* = 25 um, *dz* = 50 um and *dt* = 10^−4^ h. The parameters of the model were set to the values reported in Tables 5 and 6 (with 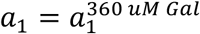), except for *a*_2_ which was set to 0.85 · 10^−9^ mL/cells h^−2^ to account for the increased killer strength of copper-induced K2 killer strains on agar plates, compared to liquid cultures. This value was calculated as follows. Upon interpolating between the last two data points in the lower-right panel of Figure 3B to estimate the value of *f_eq_* for the competition of strains K1 and K2 on agar plates, and assuming that the toxin production rate for strain K1 is unchanged with respect to liquid cultures, we found the estimate 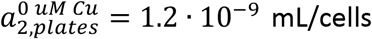 for strain K2 grown on plates with 0 uM copper and 350 uM galactose. Given that K2_b_ was a weaker killer than K2 by a factor 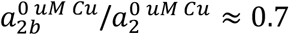 in liquid media with 0 uM copper (Table 5), we set 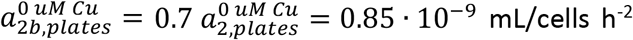. The initial condition for the radial density profiles of the two strains was set to reproduce the experiments as closely as possible. To this end, we first reproduced experimentally the initial conditions of the experiments of Figure 5 by inoculating 22 droplets of different volumes from an overnight culture of strain K1 on a lawn of strain K2_b_ using the same protocol as for the experiments of Figure 5, and droplet volumes ranging from 0.5 uL to 3 uL. We imaged the spatial distribution of cells of the two types with a fluorescence stereomicroscope at the highest magnification and analyzed the images using custom scripts written in Fiji and Mathematica to reconstruct the radial distribution of the two strains. We measured the cell densities of the two strains in the interior of the droplets, in the coffee stain rings, and outside the droplets, as well as the droplets’ radii and the width of the coffee stain rings. We integrated Equations (7–9) for each of these droplets separately (different curves in Figure 7). We have also integrated the model numerically starting from more idealized initial conditions in which strain K1 occupied the region *r ≤ r*_0_ and the strain K2_b_ occupied the region *r* ≥ *r*_0_ (i.e., the cell density of K2_b_ was set to zero for *r ≤ r*_0_), both with a density equal to the average experimental density of the experimental lawn of strain K2_b_. The dynamics of this simpler, but less accurate, simulation (Figure 7 - supplement 1) is similar to the one shown in Figure 7A-C, with a critical inoculum size of about 14 mm^2^. In the numerical integrations of the spatial model, we simulated the replica plating transfer as a dilution that preserved the relative density of the two strains at each point in space, reducing their absolute density by a factor 10^4^ and resetting the concentration of the two toxins in the agar to zero. Varying the dilution factor in the range [10^3^ – 10^s^] has almost no discernible effect on the model’s output. Numerical integrations were performed using custom Matlab scripts.

## Acknowledgements

We thank Manuel Ramírez for sending us reference K1, K2 and K^−^ strains, Elena Servienė for sending us the plasmid with the K2 DNA copy, and Manfred Schmitt and Frank Breinig for sending us the plasmid YES2.1/V5-HIS-TOPO-K1 pptox. We thank the members of the A.W.M. and D.R.N. groups for insightful comments on the research performed. A.G. thanks Marco Fumasoni for assistance in cloning. We thank Michael Laub and Michael Desai for insightful comments on the manuscript. We thank Matti Gralka and two anonymous referees for their insightful comments. A.G. was supported by the Swiss National Science Foundation, Projects P2ELP2_168498 and P400PB_180823, and work on this project was supported by the Human Frontier Science Program Grant RGP0041/2014 (to A.W.M. and D.R.N.) and NSF/Simons Center for Mathematical & Statistical Analysis of Biology at Harvard (#1764269 (NSF) and #594596 (Simons Foundation)). Work by A.G. and D.R.N. was supported by the National Science Foundation, via Grant DMR1608501 and via the Harvard Materials Science Research and Engineering Center via Grant DMR-2011754.

## Source code

Source code used for data analysis, plotting and numerical integrations of the model equations has been uploaded on GitHub at the URL: https://github.com/andreagiometto/Giometto_Nelson_Murray_2020

**Figure 3 – figure supplement 1.**
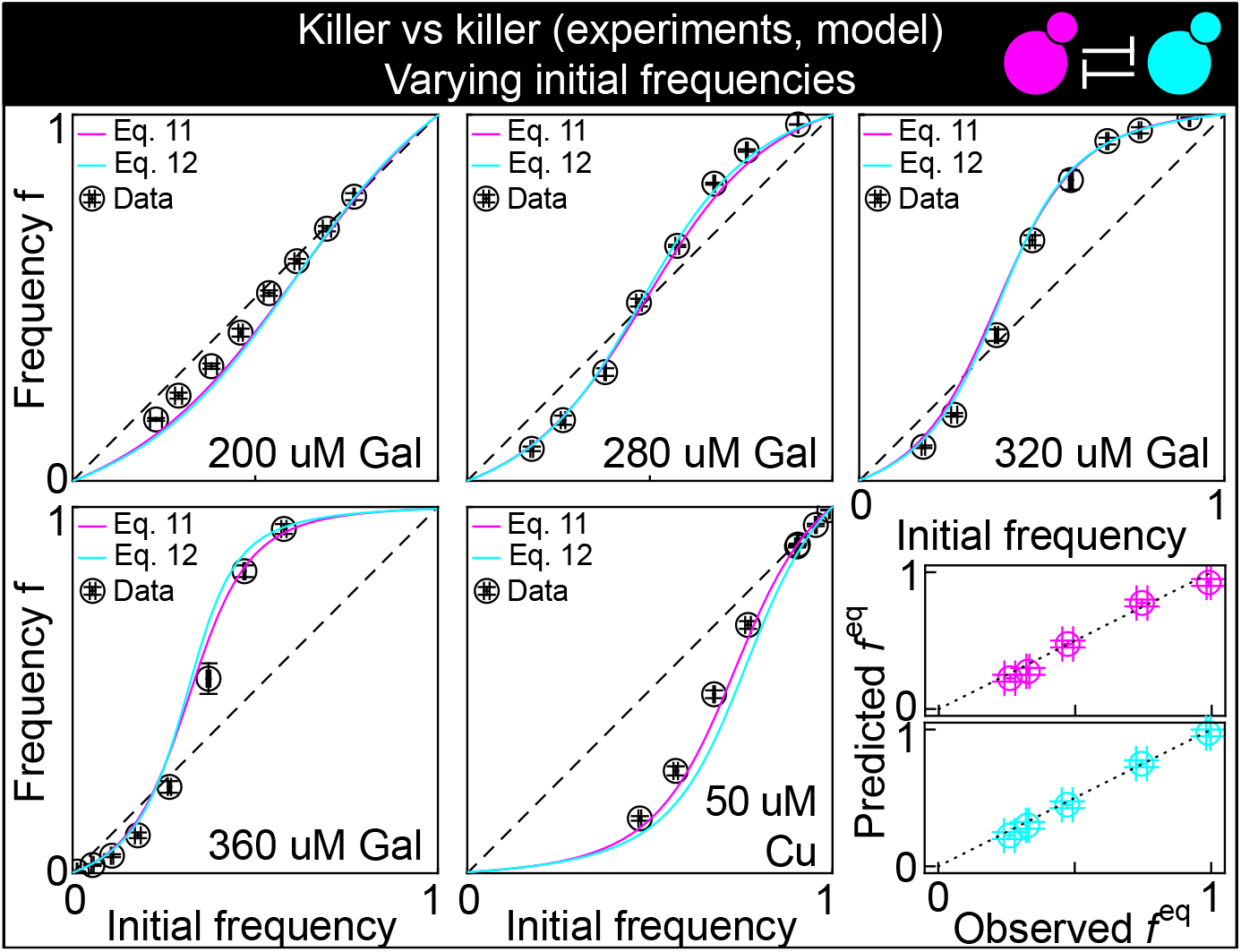
This figure shows the same data as Figure 3B, but with the predictions of the models with Equations (11) (magenta) and (12) (cyan) instead of Equation (1), modified to account for toxin production by both strains, and parametrized using competition assays between the killer strains and sensitive strains (Figure 2): K1 versus S1 and K2 versus S1. The five larger panels show the changes in the frequency of the K1 strain following a 24 h growth period, at different concentrations of the inducers and at different initial frequencies. The x axis gives the K1 frequency 14 h after inoculation, the y axis gives the K1 frequency 38 h after inoculation. The dashed and dotted lines show the 1:1 line. The critical inoculum corresponds to the intersection point between the solid and the dashed lines. The two smaller, bottom-right panels show the agreement between the predicted values of *f_eq_* (the intersection point between the solid line and the 1:1 dashed lines in the larger panels, y axis) and the measured values (the intersection point between an interpolation of the data points and the dashed 1:1 lines, x axis) for all inducer concentrations. Values using Equation (11) are shown in magenta, those for Equation (12) in cyan, with both equations modified to account for toxin production by both strains. Confidence intervals for the model predictions are not shown here because best-fit parameter errors estimated from the stationary Markov Chain distribution are very small and do not affect the model predictions significantly.

**Figure 3 – figure supplement 2.**
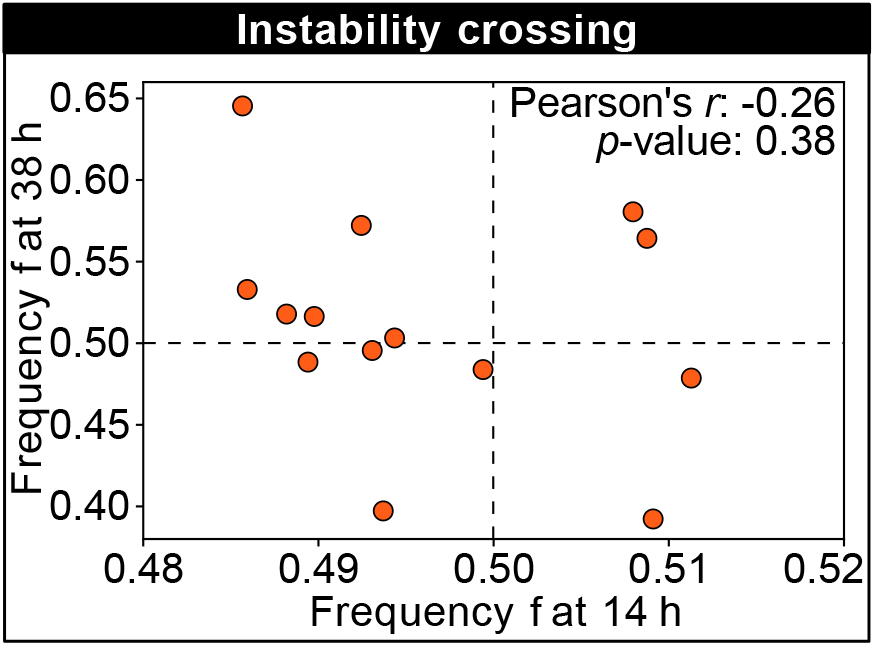
This figure shows the frequency of the K1 killer strain at the first (horizontal axis) and second (vertical axis) measurement time point for different technical replicates of a competition experiment with 250 uM galactose, which is close to the unstable equilibrium and shows replicates that tend towards different stable equilibria in the limit of large times (the same data are plotted in Figure 3D). No correlation can be detected between the frequencies at the first and second measurement time points, suggesting that the stochasticity of the dynamics is more important than the initial condition when determining which replicates will tend towards different stable equilibria in the limit of large times.

**Figure 5 – figure supplement 1.**
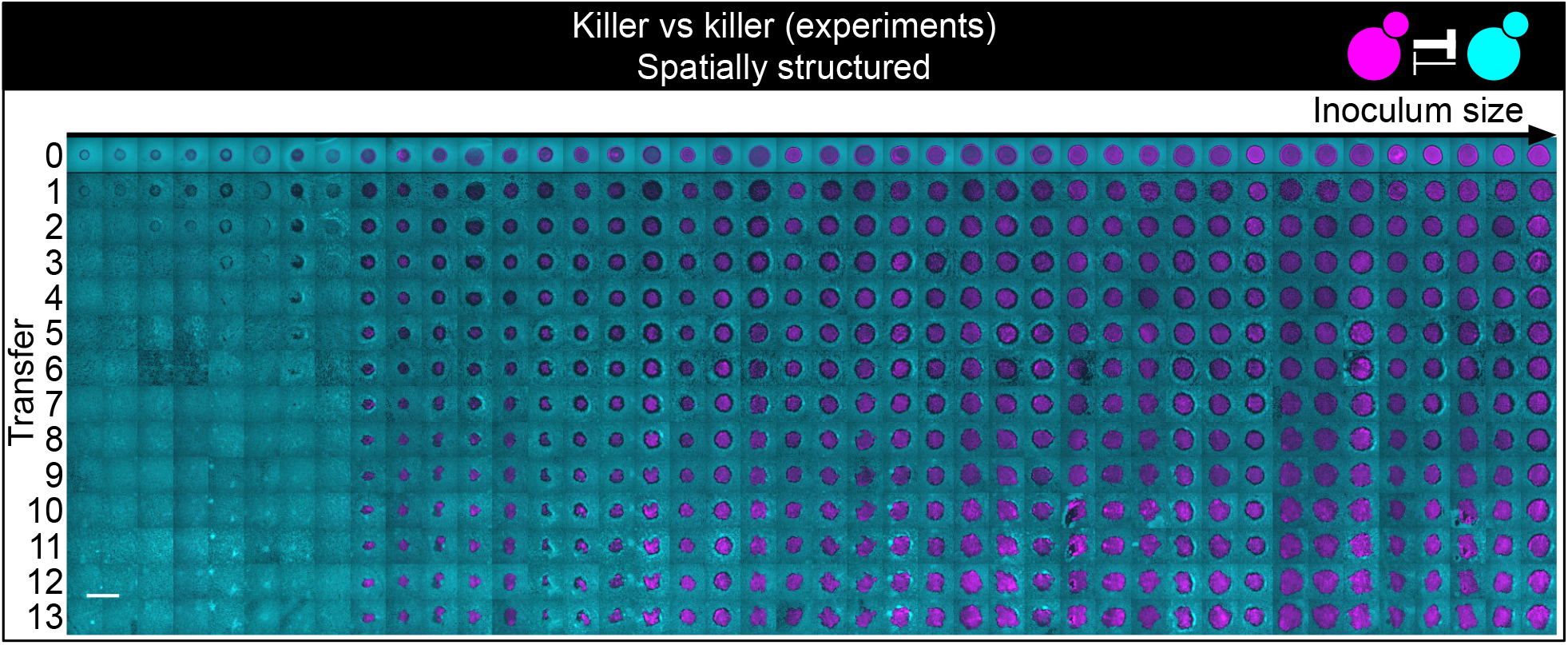
Shown here are all experimental replicates of the experiment of Figure 5. Shown from left to right are spatially structured populations of the invader K1 (magenta) and the resident K2_b_ (cyan) strains originated from depositing droplets of different volumes on a lawn of K2_b_ cells. The populations are ordered from the one with the smallest number of K1 cells (estimated via the integrated ymCherry fluorescence from strain K1) on the left to the one having the largest initial number of K1 cells at the end of the first growth cycle on the right (first row). Different rows on the same column show the same population 48 h after the previous transfer. The scale bar is 1 cm long.

**Figure 5 – figure supplement 2.**
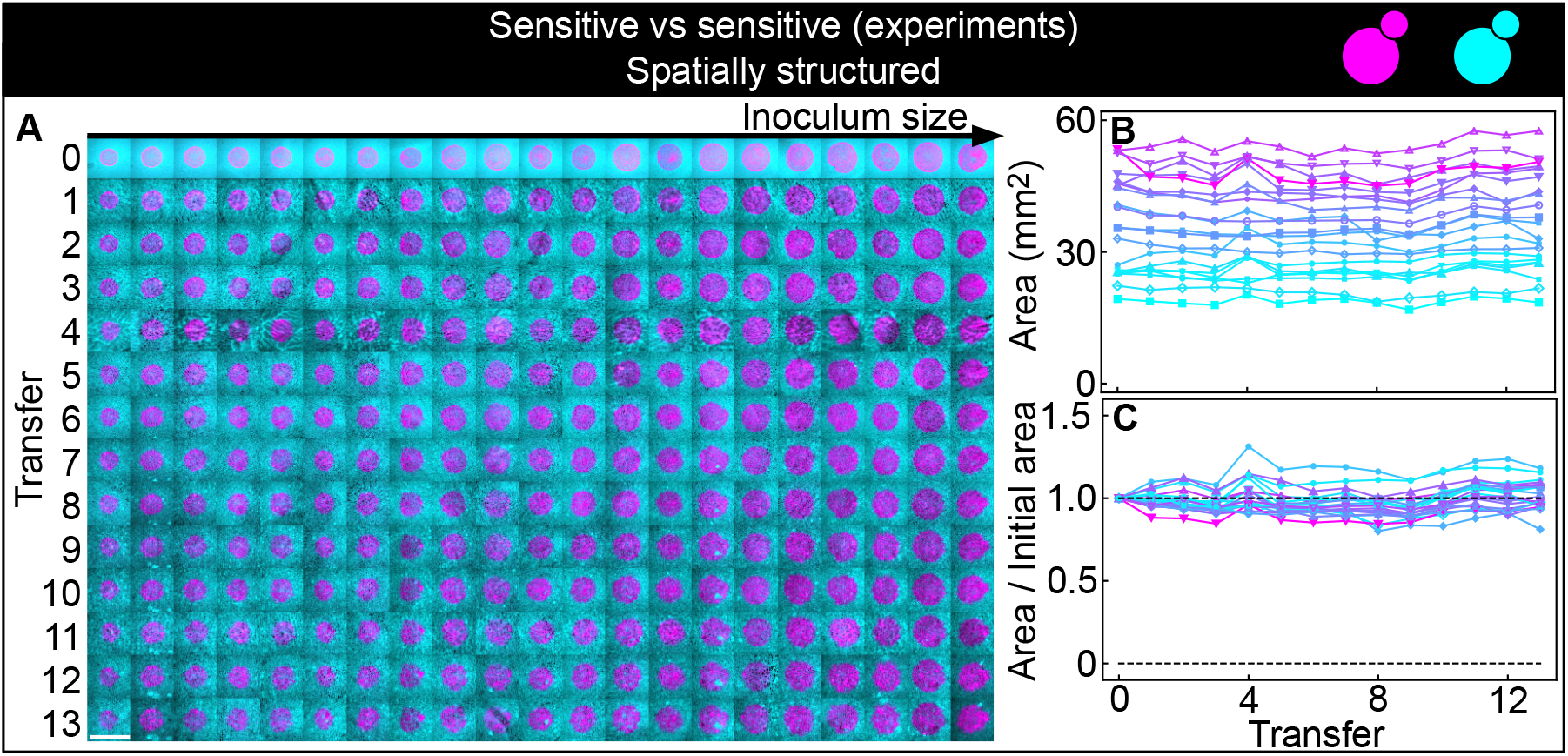
Non-antagonistic competition of sensitive, nonkiller strains in spatially structured populations. Experiments were performed in parallel to those of Figure 5, following the same experimental protocol. The strains used here carry the similar genetic constructs to those in the K1 and K2_b_ strains, but without the killer toxin genes, and thus competed solely for nutrients and for space, without direct antagonistic interactions mediated by toxins. The scale bar is 1 cm long. **(A)** Inoculations of all initial sizes maintained their shape and size throughout the experiment, following successive transfers and 48-h periods of growth. The populations are ordered based on the number of K1 cells at the end of the first growth cycle (estimated via the integrated ymCherry fluorescence of strain K1): the population with the smallest number of K1 cells is on the left, that with the largest number on the far right (first row). Different rows show the same populations after 48 h from the previous replica-plating transfer. Panels **(B-C)** show that the magenta populations maintained their areas, measured at the end of each growth period immediately before a transfer, throughout the entire experiment.

**Figure 5 – figure supplement 3.**
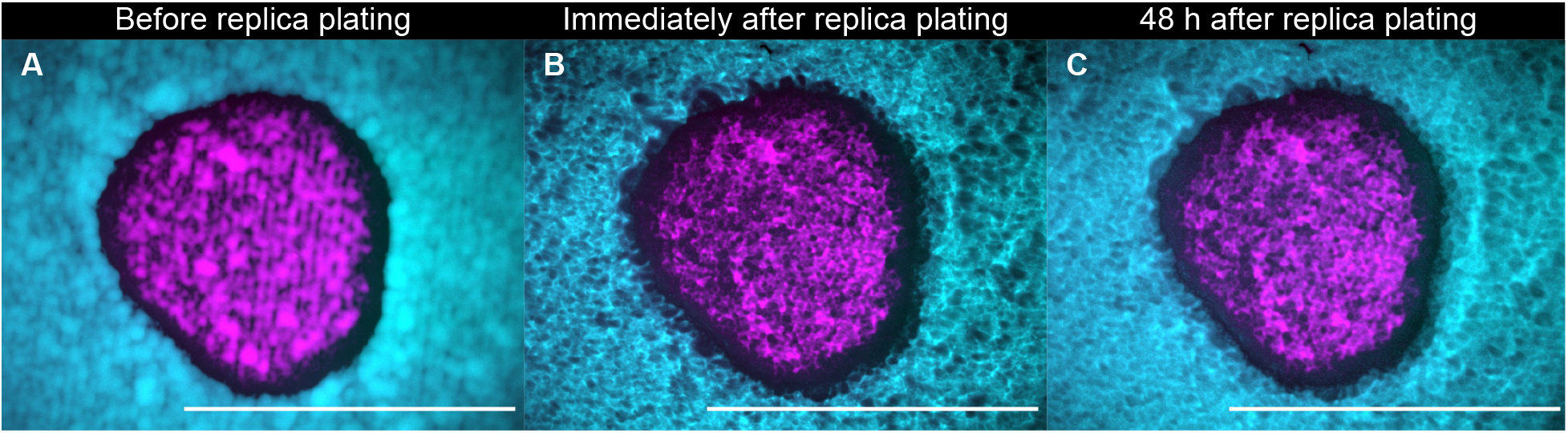
A population of K1 cells (magenta) surrounded by K2_b_ cells (cyan) imaged after 48 h of growth at 25°C immediately before replica plating (A), immediately after replica plating (B) and after further 48 h of incubation at 32°C (a temperature at which the K1 and K2 toxins are unstable) following replica plating (C). Note that panels B and C show the populations that *remain* on the plate during the replica plating transfer, and not the new, diluted copy of the populations on plates with replenished nutrients. No visible growth in the halo region can be discerned by comparing panels B and C, showing that nutrients are depleted at the end of the 48-h period of growth between transfers. Scale bars are 1 cm long. These experiments were performed using different batches of medium ingredients compared to the other experiments, and thus the activity of the two toxins and their expression rates (and thus halo widths) may differ from the other experiments.

**Figure 5 – figure supplement 4.**
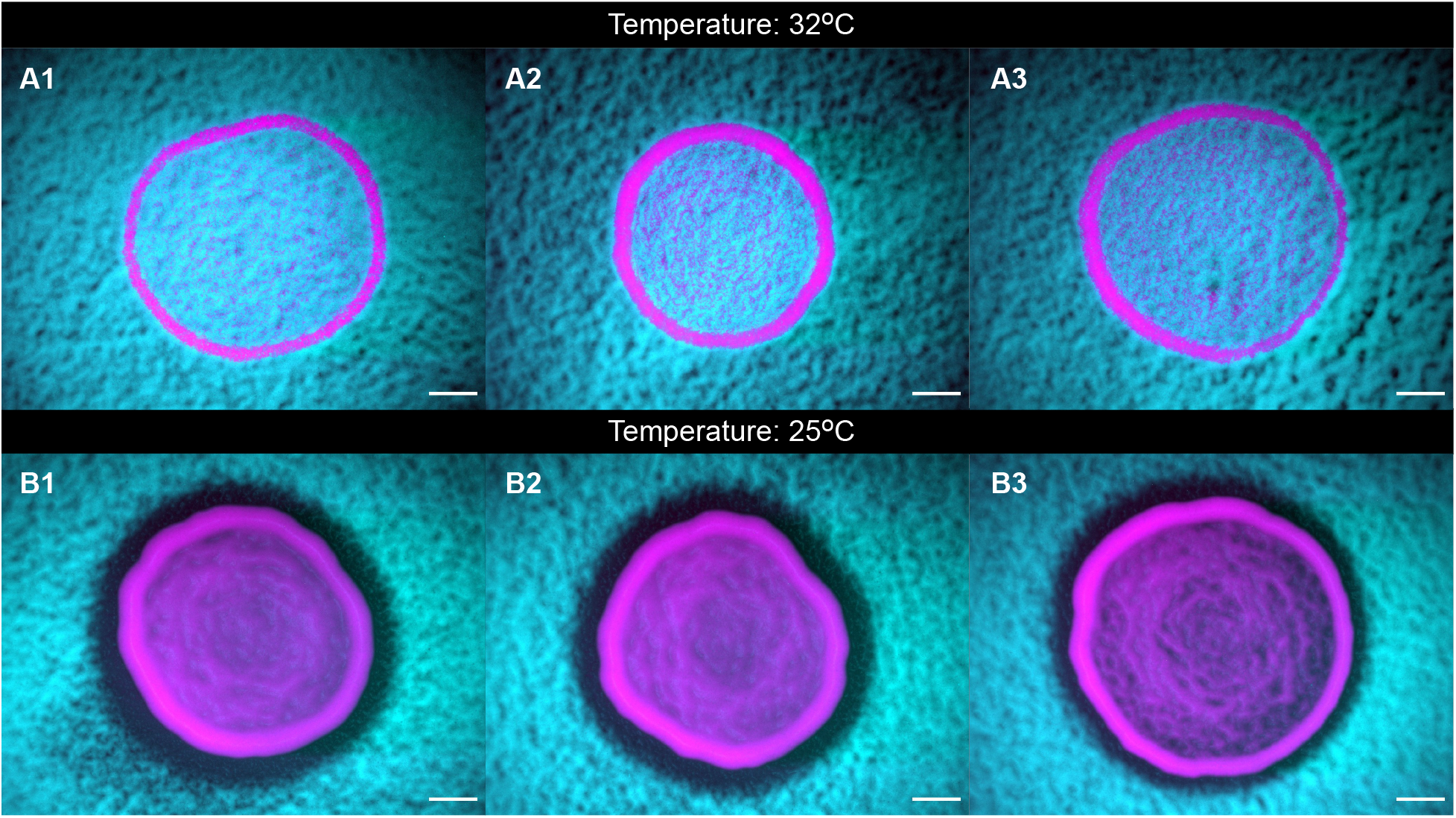
The toxins are unstable at high temperatures. Panels (A1-3) and (B1-3) show populations of strains K1 and K2_b_ grown at 25°C and at 32°C, respectively, highlighting that the toxin is unstable at 32°C, as shown by the absence of a halo at that temperature, and by the fact that the two strains coexist within the inoculum. Scale bars are 1 mm long. These experiments were performed using different batches of medium ingredients compared to the other experiments, and thus the activity of the two toxins and their expression rates (and thus halo widths) may differ from the other experiments.

**Figure 5 – figure supplement 5.**
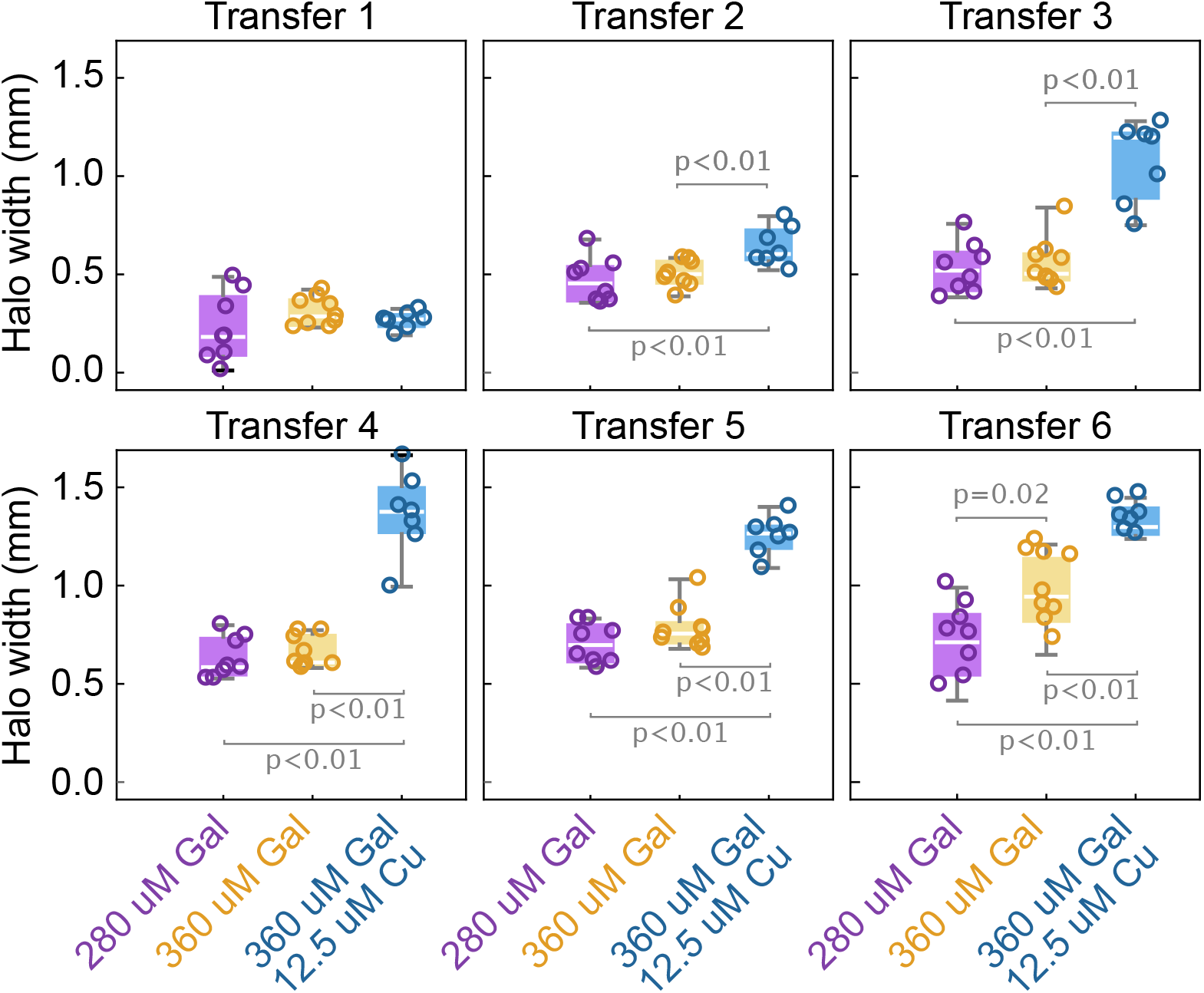
The width of the halo grows with the toxin production rate by the two strains. Box whisker plots show the width of the halo around different inoculations of the K1 strain on a lawn of K2_b_ cells, at different concentrations of the inducers (no copper was added in the 280 uM Gal and 360 uM Gal treatments), at the end of 48-h growth periods between transfers. Statistically significantly differences in the mean halo widths and the corresponding p-values are highlighted with brackets. These experiments were performed using different batches of medium ingredients compared to the other experiments, and thus the activity of the two toxins and their expression rates (and thus halo widths) may differ from the other experiments.

**Figure 6 – figure supplement 1.**
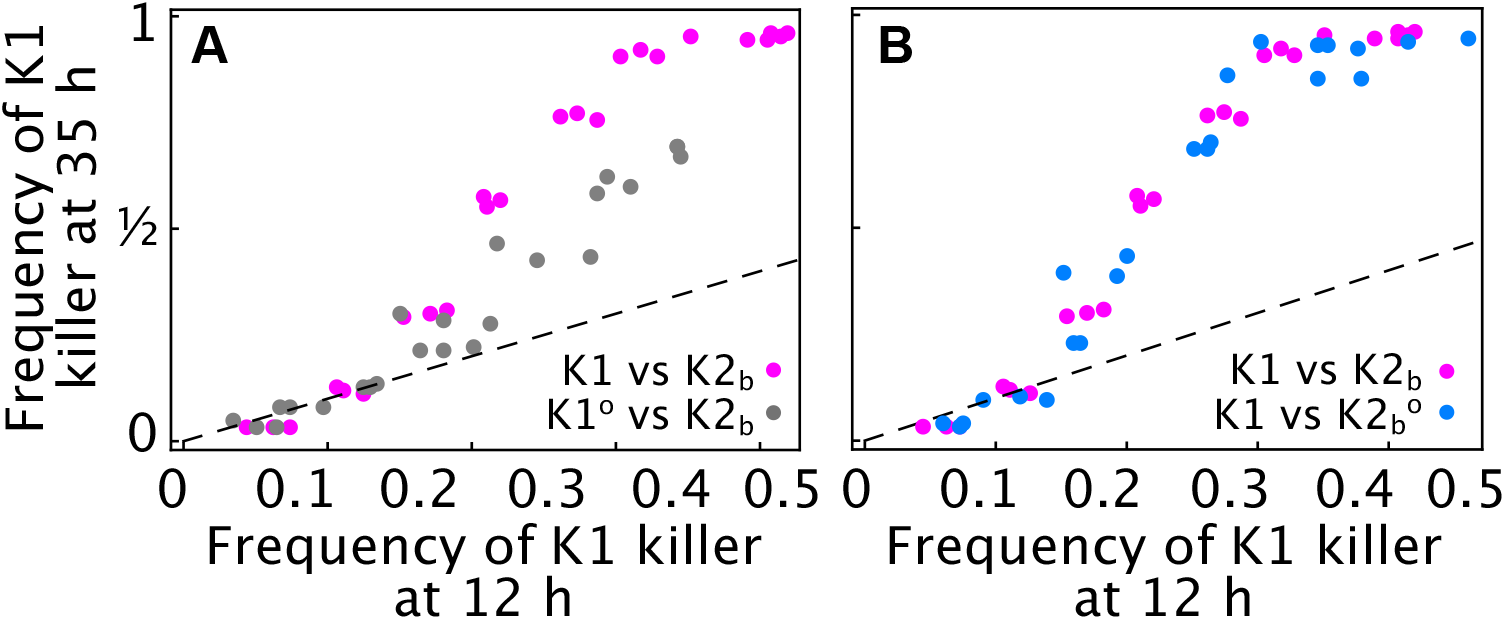
**(A)** Competition assays performed in liquid with 360 uM galactose at different initial frequencies of the K1 killer strains versus strain K2_b_ show that cells sampled from the outlier population K1^o^ (gray) increase in frequency more slowly than the ancestor K1 strain (magenta) for all initial frequencies, suggesting that K1^o^ is a weaker killer than the ancestor strain K1. The x axis gives the K1 frequency 12 h after inoculation, the y axis gives the K1 frequency 35 h after inoculation. **(B)** Competition assays performed in liquid with 360 uM galactose at different initial frequencies of the killer strain K1 versus strain K2_b_ (magenta) and K2_b_^o^ (K2 cells sampled from the region around the outlier population K1^o^, light blue) show that K2_b_ and K2_b_^o^ cells have comparable killer strengths, given that their frequency falls (seen as a rise in the frequency of strain K1) at similar rates. Different dots show measurements from different technical replicates. Dashed lines are 1:1 lines.

**Figure 6 – figure supplement 2.**
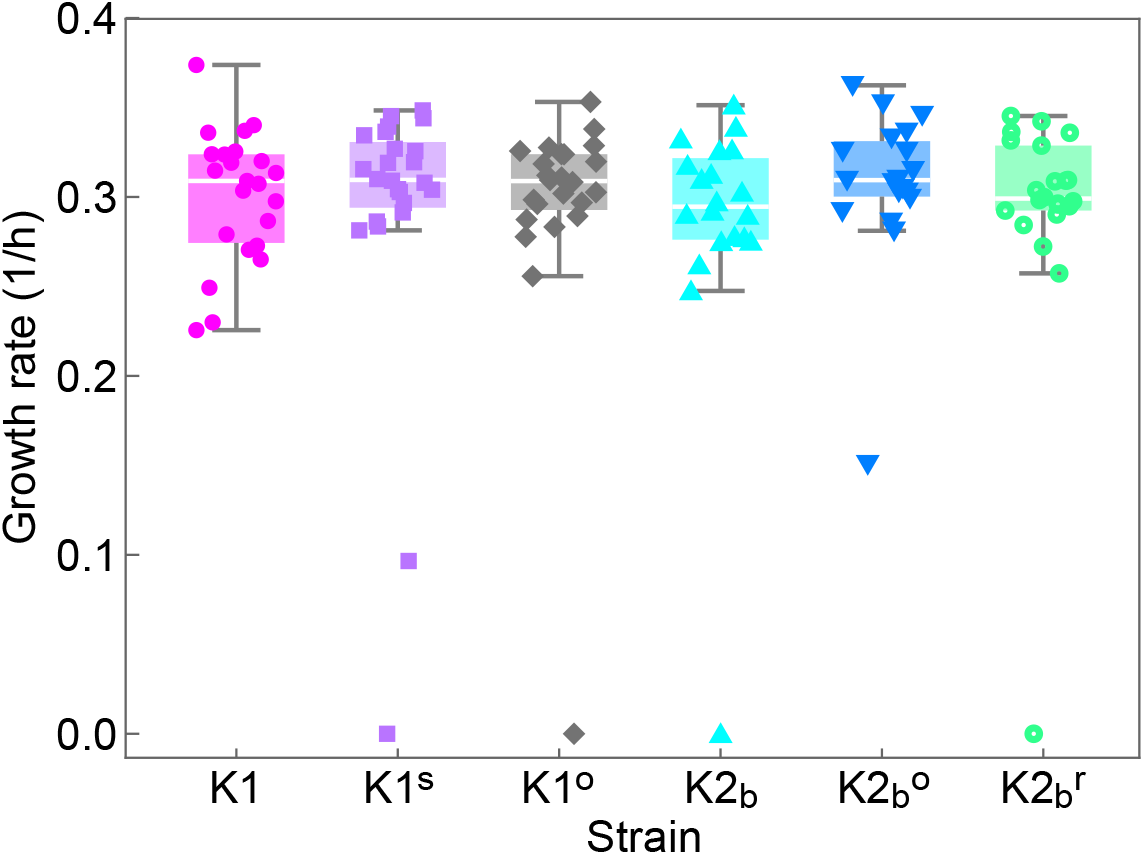
Single-cell growth rates of the ancestor strains (K1, K2_b_) and of cells isolated at the end of the experiment of Figures 5 and 6 (K1^s^, K1^o^, K2_b_^o^, K2_b_^r^) grown on YPD agar plates identical to the ones used for the experiments of Figures 5 and 6, but in the absence of inducers. Droplets of each strain were deposited on a plate on a grid at distances of 1 cm from each other and the location of each strain within the grid was randomized. Cells on the plate were imaged using an inverted microscope with 50X objective every 20 min for 6 h, during which the plate was kept at 25°C using a stage-top incubator. During the 6 h, cells formed micro-colonies of up to 32 cells, except for a few non-dividing or very slow dividing cells that we excluded from the analysis, but whose growth rates are shown in the plot. Data points show growth rates obtained by fitting exponential growth curves to experimental data for the number of cells in individual micro-colonies. T-tests performed between all pairs of strains/mutants gave no statistically significant differences between their growth rates (all corresponding p-values are larger than 0.05). Box whiskers plots show the median growth rate (white horizontal lines), the 25% and 75% percentiles (colored boxes) and the maximum and minimum growth rate for each strain (whiskers). Slow- and non-growing cells treated here as outliers have not been included in the computation of medians, percentiles, and minima/maxima.

**Figure 6 – figure supplement 3.**
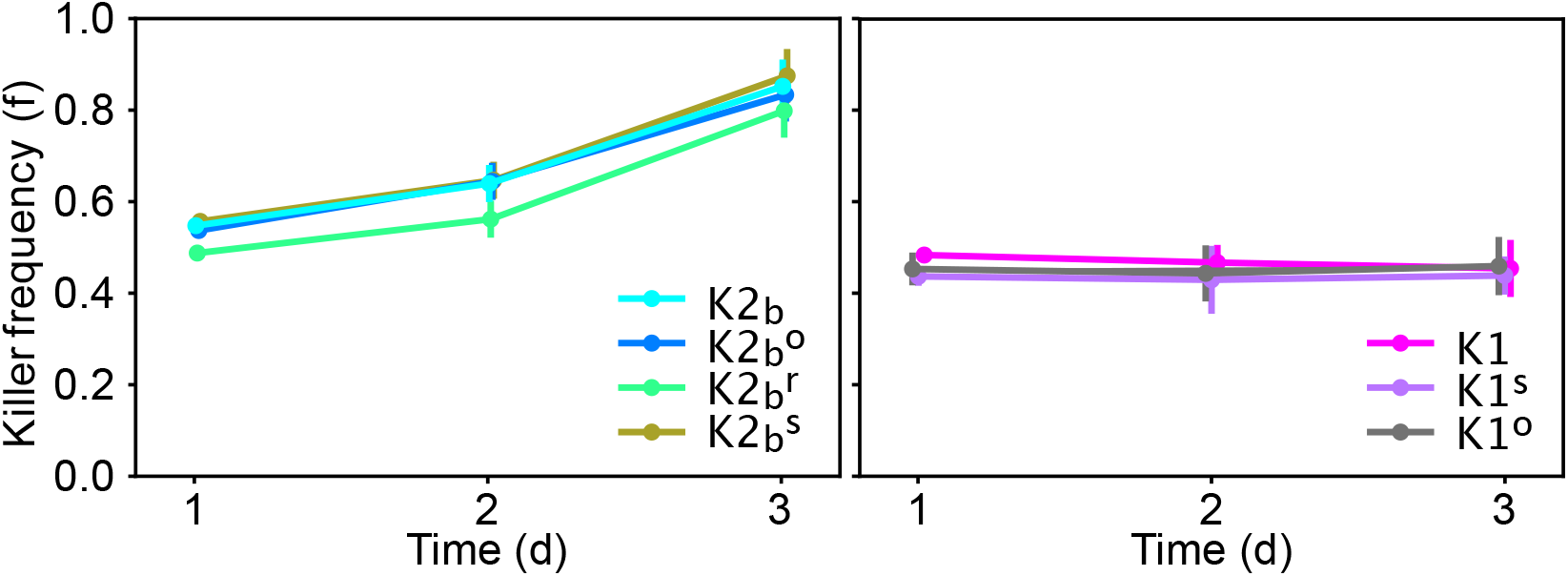
Relative frequency of the ancestor strains (K1, K2_b_) and of cells isolated at the end of the experiment of Figures 5 and 6 (K1^s^, K1^o^, K2_b_^o^, K2_b_^r^) in competition against sensitive strains in well-mixed liquid cultures diluted daily in YPD medium, in the absence of inducers. Data points show the relative frequency of killer strains across 12 technical replicates, error bars are standard deviations. The left panel shows competitions between cells with the K2 killer toxin gene against the sensitive strain S1, the right panel shows competitions between strains with the K1 killer toxin gene against the sensitive strain S2. Experiments were initialized with a 50-50 mixture of two overnight cultures of each strain and were measured after 1 d, 2 d and 3 d from inoculation, right before dilution. Mixed competition assays performed with strains carrying the K1 killer toxin gene induced by the promoter *P_GAL1_* (strains K1, K1^s^ and K1^o^) versus strain S2 can be used to directly detect differences in reproductive fitness, because no toxin is produced in the absence of inducer (galactose) and thus the relative frequency of the two strains varies due to differences in reproductive fitness alone. Mixed competition assays between strains carrying the K2 killer toxin gene induced by the promoter *P_CUP1_* (K2_b_, K2_b_^r^ and K2_b_^o^) versus strain S1, instead, can only detect the joint effect of differences in reproductive fitness and killer activity, given that the promoter *P_CUP1_* has non-zero expression even in the absence of inducer (copper). T-tests performed between all pairs of competitions between killer strains/mutants of the same type (i.e., expressing either the K1 or K2 toxin) and the corresponding sensitive strains (figure below) give no statistically significant differences between the rates at which the relative frequency of killer strains in competition assays vary with time (all corresponding p-values are larger than 0.05). These experiments were performed using different batches of medium ingredients compared to the other experiments, and thus the activity of the two toxins and their expression rates may differ from the other experiments.

**Figure 7 – figure supplement 1.**
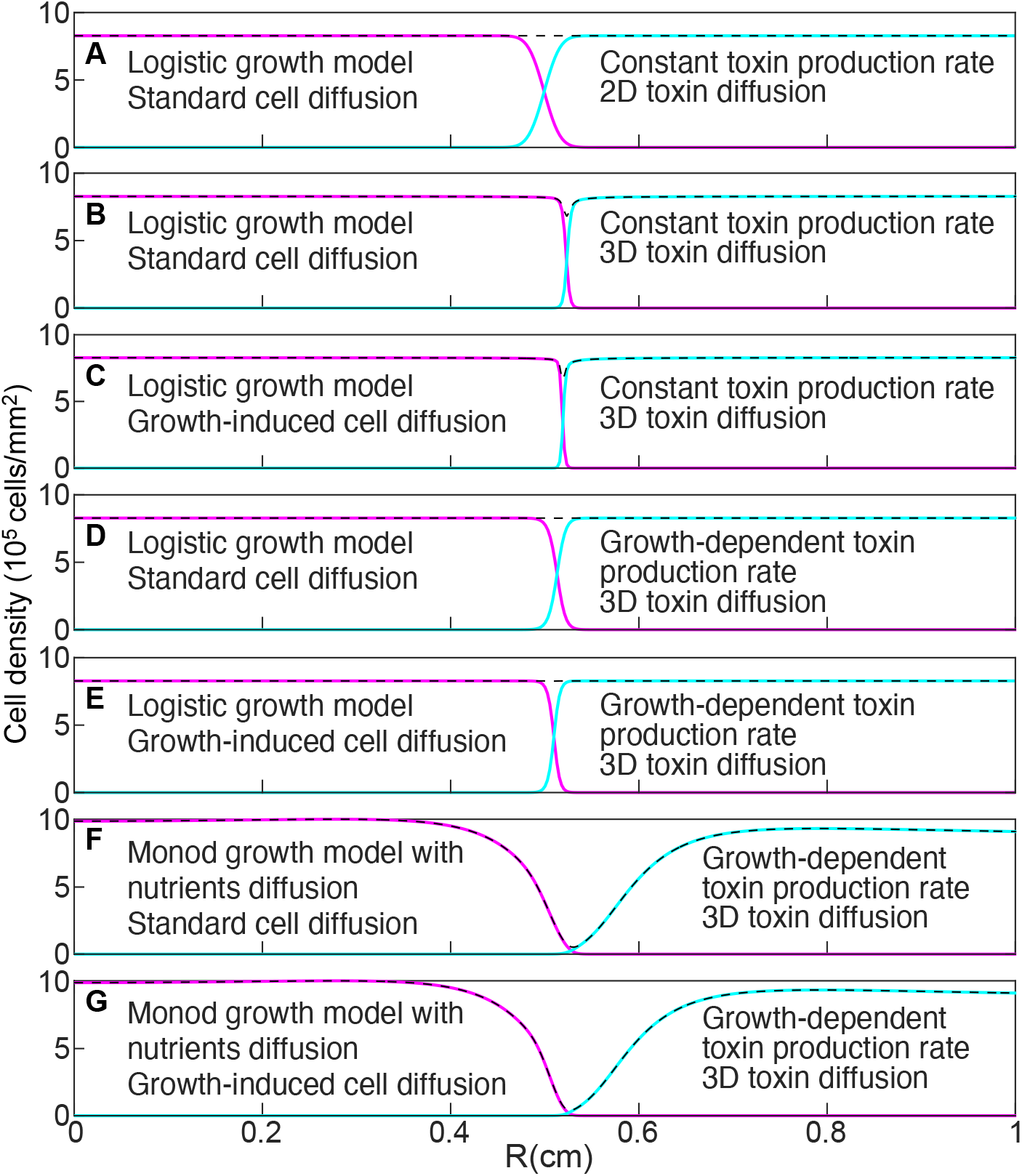
Models that do not account for nutrient diffusion fail to reproduce the formation of the halo between two antagonist strains. Magenta and cyan curves show the density profiles of K1 and K2 killer strains. The dashed, black lines show the total cell density and are the sum of the magenta and cyan density profiles. All models were parametrized using competition experiments between killer and sensitive strains, and parameters corresponding to strains K1 and K2_b_ competing with 360 uM galactose and 0 uM copper were used. In the initial conditions strain K1 occupied the entire region *r ≤r*_0_ and strain K2_b_ occupied the region *r > r*_0_, with *r*_0_ = 5 mm. The figures show density profiles after six transfers. The model in panel A corresponds to Equations 6. The model in panel G corresponds to Equations 7–9. The characteristic features of intermediate models are reported in each panel.

**Figure 7 – figure supplement 2.**
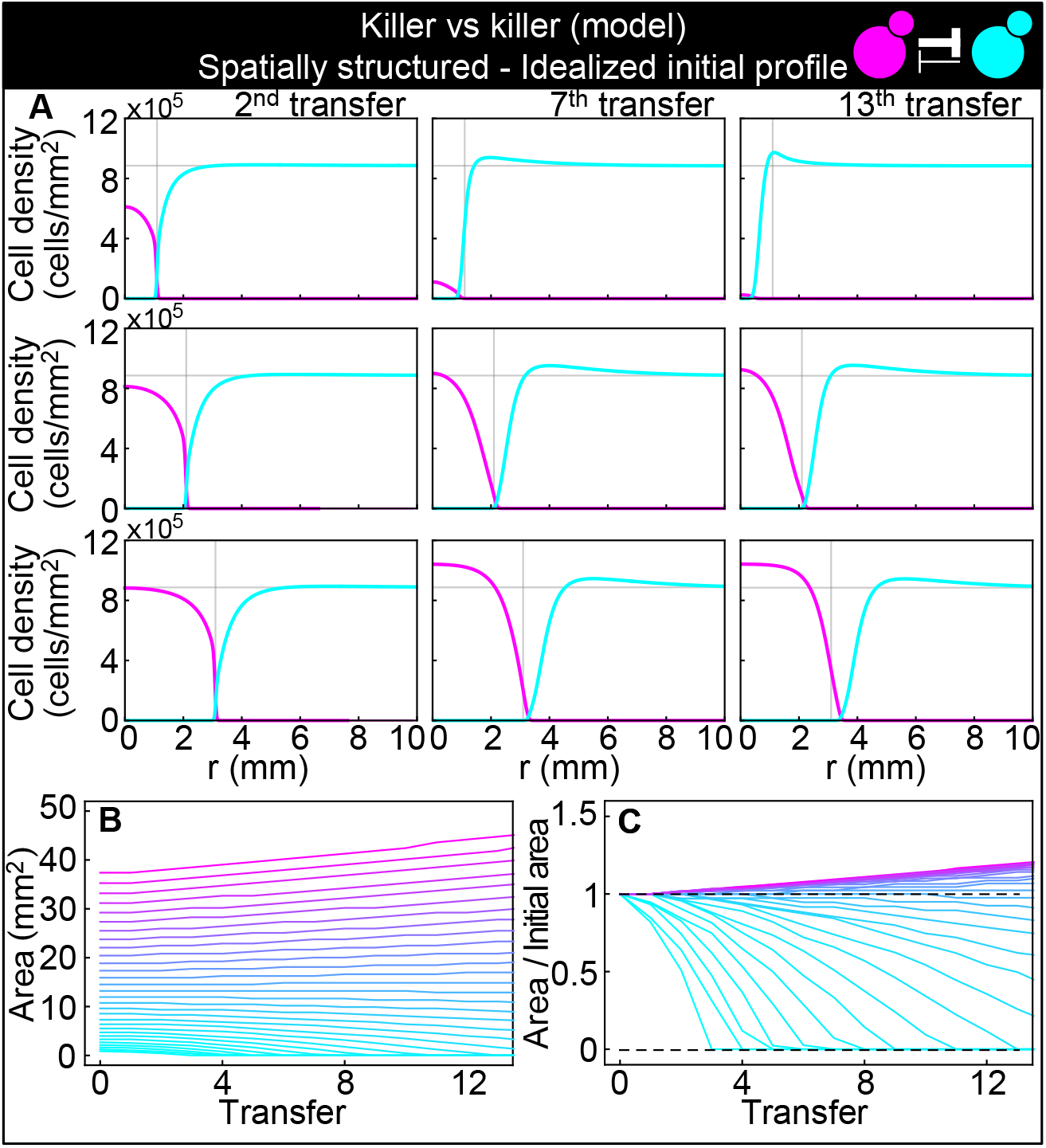
Numerical integrations of the spatial model (Equations 7–9), parametrized using the Experiments of Figure 2 and other values taken from the literature, can qualitatively reproduce the experimental dynamics of antagonistic competition between strains K1 and K2_b_ in spatially structured populations. Same plots as Figure 7, but with idealized initial conditions in which strain K1 occupied the entire region *r ≤ r*_0_ and strain K2_b_ occupied the region *r > r*_0_, where the initial radius *r*_0_ was varied between 5 and 34 mm. **(A)** Simulated K1 inoculations smaller than the critical inoculum (first row) expanding on a landscape occupied by strain K2_b_ fail to establish and expand, whereas larger inoculations do (second and third row). The model reproduces the formation of the halo, a region without cells at the interface between the two strains (second and third row). **(B)** Area covered by each simulated K1 population expanding on a landscape occupied by strain K2_b_ at the end of each 48-h growth period between simulated transfers, color coded from cyan to magenta according to the total K1 population of each replica at the end of the first growth period. **(C)** Same data as in (D), divided by the initial area to highlight relative changes. Comparison with Figure 7B and 7C shows that the presence of K2 cells within the K1 inoculum leads to faster extinction of K1 cells in inocula below the critical size, is required for the initial retreat of K1 populations above the critical size, and leads to a more abrupt transition between the dynamics of K1 populations that go extinct and those that thrive.

**Figure 7 – figure supplement 3.**
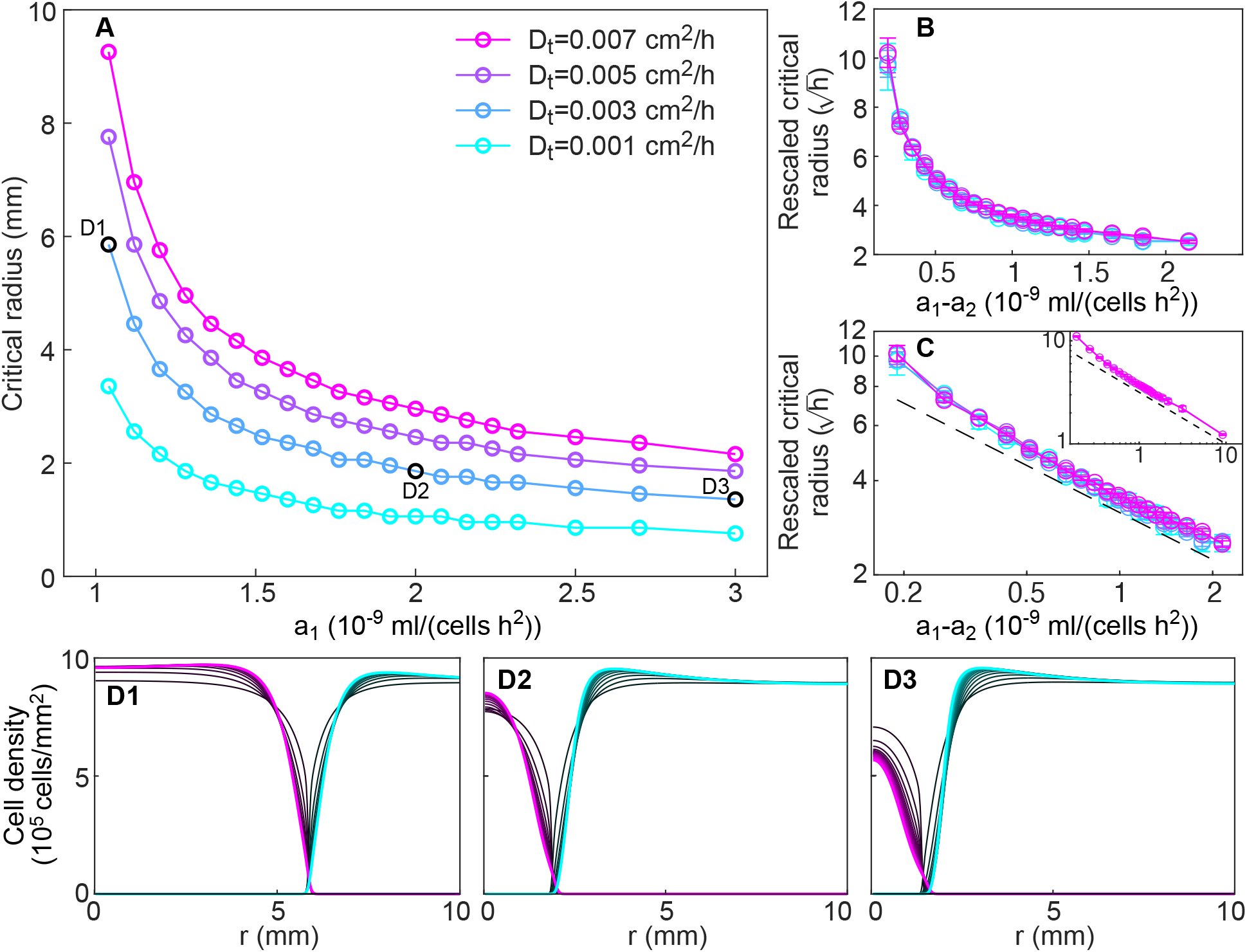
**(A)** Size of the critical radius for different values of the toxin production rate of the invader strain *a*_1_ and of the toxin diffusion rate (different curves), for a fixed value of the toxin production rate of the resident strain, *a*_2_ = 0.85 × 10^−9^ ml/(cells h^2^) in numerical simulations of the model (Equations 8–9), starting from idealized initial conditions as for Figure 7 - supplement 2. Critical radii were computed by integrating Equations 8–9 over 20 consecutive transfers for different values of the inoculum radius (varied with step size 0.1 mm), measuring the total number of K1 cells at the end of each transfer, and identifying the largest and lowest values of the inoculum radius such that the total number of K1 cells was decreasing and increasing, respectively, in the last five transfers. Panels **(B-C)** shows that curves corresponding to different values of the toxin diffusion coefficient *D_t_* collapse onto a single curve when dividing the critical radius by a factor 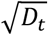. In panel B, rescaled critical radii and *a*_1_ − *a*_2_ are plotted in linear scale. In panel C, rescaled critical radii are plotted in log-log scale versus the difference between the invader and resident strain toxin production rates *a*_1_ − *a*_2_, to highlight the approximate power-law dependence for large values of *a*_1_ − *a*_2_ (the dashed black line is proportional to 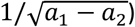. The inset of panel C shows that the power-law behavior extends to *a*_1_ − *a*_2_ = 10^8^ ml/(cells h^2^) (only data for *D_t_* = 0.005 cm^2^/h and *D_t_* = 0.007 cm^2^/h are shown because smaller values of *D_t_* would require a smaller radial integration step). **(D1-D3)** Density profiles of the two strains at the end of each growth period between transfers (black curves), starting from uniform spatial distributions of the invader strain within the critical radius and of the resident strain outside the critical radius. Shown in magenta and cyan are the stationary density profiles of the two strains in the limit of large times, after the profiles have reached stationarity.

**Figure 7 – figure supplement 4.**
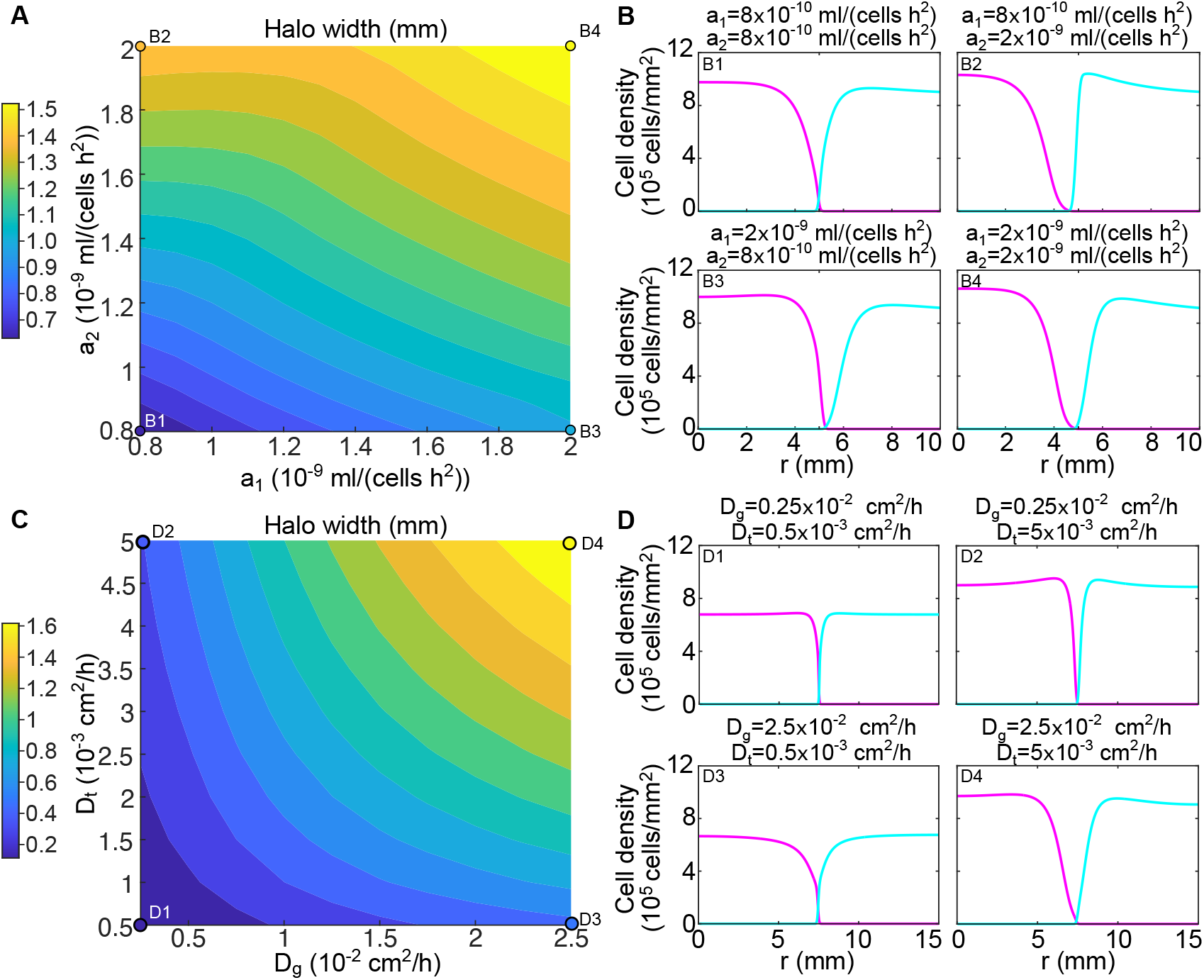
The width of the halo and the shape of the two strains’ density profiles varies with the parameters of the model. **(A)** Width of the halo region between the invader and the resident strain for different values of the toxin production rates *a*_1_ and *a*_2_ in numerical simulations of the model (Equations 8–9). The halo width was measured as the distance between the radius at which the resident density profile reaches half its maximum and the radius at which the invader density profile reaches half its maximum. **(B)** Density profiles of the two strains for different values of the toxin production rates, corresponding to the points highlighted in panel A. **(C)** Width of the halo region between the invader and the resident strain for different values of the toxin diffusion rate *D_t_* and the glucose diffusion rate *D_g_* in numerical simulations of the model (Equations 8–9), with the toxin production rates set to *a*_1_ = *a*_2_ = 1.5 · 10^−9^ ml/(cells h^2^). **(D)** Density profiles of the two strains for different values of the toxin and glucose diffusion rates, corresponding to the points highlighted in panel C. In all the panels, the populations were diluted every 48 h and the halo width was measured after five dilutions. In the initial conditions strain K1 occupied the entire region *r* ≤ *r*_0_ and strain K2_b_ occupied the region *r > r*_0_, with *r*_0_ = 0.5 cm.

